# Nucleolin is essential for rabbit hemorrhagic disease virus replication by providing a physical link in replication complex formation

**DOI:** 10.1101/2020.05.13.094185

**Authors:** Jie Zhu, Qiuhong Miao, Hongyuan Guo, Ruibin Qi, Aoxing Tang, Dandan Dong, Jingyu Tang, Guangzhi Tong, Guangqing Liu

## Abstract

Rabbit hemorrhagic disease virus (RHDV) is an important member of the *Caliciviridae* family and cannot be propagated *in vitro*, which has greatly impeded progress of investigating its replication mechanism. Construction of an RHDV replicon system has recently provided a platform for exploring RHDV replication in host cells. Here, aided by this replicon system and using two-step affinity purification, we purified the RHDV replicase and identified its associated host factors. We identified rabbit nucleolin (NCL) as a physical link required for the formation of RHDV replication complexes (RCs), by mediating the interaction between other host proteins and the viral RNA replicase, RNA-dependent RNA polymerase (RdRp). We found that RHDV RdRp uses an amino acid (aa) region spanning residues 448–478 to directly interact with NCL’s RNA-recognition motif 2. We also found that the viral p16 protein uses a highly conserved region (^35^Cys–Ile–Arg–Ala^38^ or CIRA motif) to specifically bind the N-terminal region of NCL (aa 1–110) and that RHDV p23 uses a specific domain (aa 90–145) to bind NCL’s RNA-recognition motif 1. Disrupting these protein–protein interactions severely weakened viral replication. Furthermore, NCL overexpression or knockdown significantly increased or severely impaired, respectively, RHDV replication. Collectively, these results indicate that the host protein NCL is essential for RHDV replication and plays a key role in the formation of RHDV RCs. The mechanisms by which NCL promotes viral replicase assembly reported here shed light on viral RC biogenesis and may inform antiviral therapies.

**Author summary:** Rabbit hemorrhagic disease virus (RHDV) is the causative agent of highly contagious and lethal hemorrhagic disease in the European rabbit, but the host factors involved in RHDV replication remain poorly understood. In the present study, we isolated RHDV replication complex (RC) for the first time and identified its main components. We found that nucleolin (NCL) plays a key role in the formation of the RHDV RC. NCL not only interacts with viral replicase (RdRp), it also specifically binds to other important host factors. In addition, we proved that NCL is necessary for RHDV replication because the level of RHDV replication is significantly affected by knocking down the NCL gene in cells. Together, our data suggest that RHDV completes its replication by hijacking NCL to recruit other viral proteins and host factors, thereby assembling the RC of RHDV.

## Introduction

Rabbit hemorrhagic disease virus (RHDV) is the causative agent of rabbit hemorrhagic disease (RHD), which primarily infects the wild and domestic European rabbit (*Orcytolagus cuniculus*) [1] and is characterized by liver degeneration, diffuse hemorrhaging and high mortality [2,3]. However, the molecular mechanisms responsible for RHDV replication remain poorly understood, mainly due to the lack of a robust cell culture system for propagation of the virus.

RHDV is a nonenveloped positive-sense single-stranded RNA virus, which belongs to the family *Caliciviridae*, genus *Lagovirus*. RHDV virions contain the genomic RNA (gRNA) and an additional 2.2 kb of subgenomic RNA (sgRNA), which is collinear with the 3′ end of the gRNA [1]. The gRNA of RHDV consists of a positive-sense single-stranded molecule of 7,437 nucleotides with a virus-encoded protein, VPg, which is covalently attached to its 5′ end [4,5]. The gRNA also contains two slightly overlapping open reading frames (ORFs) of 7 kb (ORF1) and 351 nucleotides (ORF2). ORF1 is translated into a large polyprotein that is cleaved into the major structural protein VP60, the capsid protein, and seven nonstructural proteins: p16, p23, helicase, p29, VPg, protease, and RNA-dependent RNA polymerase (RdRp). ORF2 encodes the minor structural protein VP10 [6-8]. The sgRNA, which only encodes VP60 and VP10, usually contributes to the production of high levels of products required during intermediate and late stages of infection [9]. Flanking the coding regions of RHDV is a 5′ terminal noncoding region of nine nucleotides and a 3′ terminal noncoding region of 59 nucleotides [8].

The function of some of the nonstructural proteins encoded by the genome of caliciviruses has been identified and/or predicted by relying on previous knowledge gathered from members of the closely related *Picornaviridae* family. For RHDV, two nonstructural proteins might be involved in the replication of viral RNA, a helicase and an RdRp, and a protease is involved in the autoproteolytic processing of the large viral polyprotein encoded by ORF1 [7,10,11]. The genome-binding protein VPg is covalently linked to both genomic and subgenomic RHDV RNAs at the 5’ end. Our previous study suggested that the VPg protein serves as a novel cap substitute during the initiation of RHDV translation [5]; however, the precise function of the RHDV nonstructural proteins (p16, p23, and p29) remains unclear.

Following attachment of RHDV to the cell surface, internalization and desencapsidation occur, leading to release of the viral genome into the cytoplasm of the host cell. The virus life cycle then proceeds to translation of the polyprotein precursor encoded by the viral genome through interaction with the host cellular machinery. The gRNA and the sgRNA covalently-linked VPg use the cellular translation machinery, positioning the ribosome at the start codon AUG without ribosome scanning and initiating translation. Posttranslational proteolytic processing by the viral gRNA encoded protease cleaves the polyprotein precursor into the mature nonstructural proteins and, in RHDV, into the capsid protein VP60. The nonstructural proteins, helicase and RdRp, then form a replication complex (RC), synthesizing complementary negative-sense RNA from the gRNA, which is used as a template for the synthesis of gRNA and sgRNA.

The positive-strand RNA viruses share a conserved replication mechanism in which viral proteins induce host cell membrane modification to assemble membrane-associated viral RC [12]. Viruses hijack host factors to facilitate this energy-consuming process [13]. However, understanding the detailed molecular mechanism of how viral proteins hijack host factors for RC assembly has been hampered by the lack of a suitable *in vitro* culture system for RHDV. Identification of replicase-associated host factors and dissection of their roles in RC assembly will shed light on the molecular mechanism of RHDV replication. In 2013, we developed an RHDV replicon system, which has the ability to automatically replicate in RK-13 cells [14]. Construction of this RHDV replicon system has provided a platform for exploring RHDV replication in host cells. In 2017, we successfully constructed mutant RHDV (mRHDV) in RK-13 cells *in vitro*, which has a specific receptor-recognition motif (Arg-Gly-Asp) on the surface of the capsid protein that is characterized by two aa substitutions. mRHDV is recognized by the intrinsic membrane receptor (integrin ɑ3β1) of RK-13 cells, by which mRHDV gains entry, replicates, and imparts apparent cytopathic effects [15].

In this study, an RHDV RC was isolated for the first time and its main components were identified. We found that nucleolin (NCL) plays a key role in the formation of viral RCs. Our data showed that NCL is a link between viral replicase and host proteins. In addition, we demonstrated that NCL is necessary for RHDV replication because the replication level of RHDV is significantly affected by knocking down the *NCL* gene in cells.

## Results

### Tagging of RHDV replicase (RdRp) in the context of a viral replicon

To discover the host factors that are involved in RHDV replication, we attempted to purify the viral RCs formed during viral replication and identify the associated host factors. Previously, the researchers successfully identified hepatitis C virus (HCV) replicase-associated RC components by inserting His and HA tags into the HCV replicon replicase NS5A and NS5B (RdRp) for affinity purification [16]. Here, we aimed to affinity tag RdRp with two different tags to facilitate tandem affinity purification. We generated a recombinant replicon by introducing a His or HA tag into RdRp (aa sites: 25, 82, 442, or 483, respectively) of the RHDV replicon (Fig. 1A). Moreover, as predicated using the SWISS-MODEL online tool (https://swissmodel.expasy.org/), we found that insertion of the His and/or HA tag into these sites would have no effect on the structure of RdRp (Fig. S1). Fluc activity analysis showed that RHDV-luc-His_25_, RHDV-luc-HA_442_, and RHDV-luc-HA_483_ replicated similarly to the untagged RHDV replicon, whereas the replication ability of RHDV-luc-His_82_ was significantly inhibited in RK-13 cells (Fig. 1B). The same results were obtained in immunoblotting (IB) detection (Fig. 1C). The luciferase activity from the replicon lacking the RdRp gene has previously been shown to be approximately 4-logs lower than that of the wild type replicon [14]. In RHDV replicon, the viral sequence was generated as a consequence of polymerase II transcription from the cytomegalovirus (CMV) promoter, and the authentic 3’ end of the viral genome was under controlled by a hepatitis delta virus ribozyme [17,18]. Subsequently, RdRp, VPg, and other non-structural proteins were translated in a cap-dependent manner by the host cell. The luciferase is expressed from the subgenomic RNA which generated by the RHDV RdRp. Thus, we could confirm insertion of His or HA tag at aa sites 25, 442 or 483 of RdRp.

**Figure 1.**
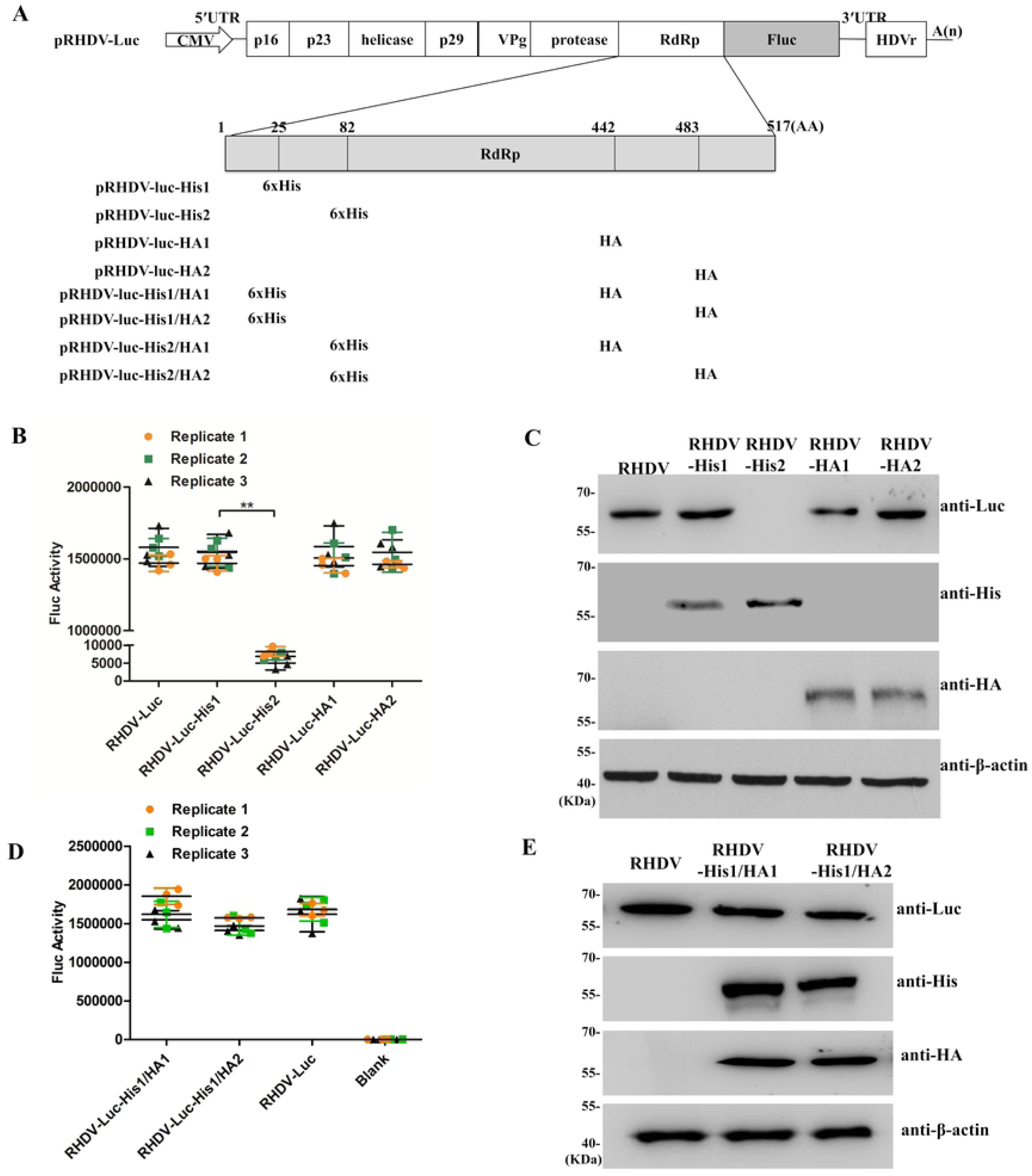
Tagging of RHDV RdRp in the viral replicon. **(A)** Schematic of RHDV-His/HA replicon constructs. The HA and 6×His peptide sequences are inserted into the RdRp aa sequence at 25, 82, 442, or 483 sites, respectively. **(B, D)** Effect of the inserted tag on viral replicon activity. RK-13 cells were transfected with recombinant RHDV replicons. Luciferase activity in cell lysates was measured at 48 hpt. Student *t*-tests and analysis of variance were used for statistical analyses. \**p* < 0.05 and \*\**p* < 0.01. The number of cells used in all replicate experiments was similar. **(C, E)** Western blotting of recombinant RHDV replicons in RK-13 cells with the antibodies indicated. β-actin was used as an internal control.

Subsequently, we introduced HA and His peptides into RdRp simultaneously to obtain a double-affinity-tagged replicon (Fig. 1A). Fluc activity and IB analyses showed that RHDV-luc-His_25_/HA_442_ and RHDV-luc-His_25_/HA_483_ replicated similarly to the untagged RHDV replicon in RK-13 cells (Fig. 1D and 1E). Therefore, we used RHDV-luc-His_25_/HA_442_ in affinity purification assays.

### Identification of host factors associated with RHDV replicase

After transfection of RK-13 cells with the RHDV-luc-His_25_/HA_442_ replicon for 48 h, solubilized cell lysates were sequentially purified using the HA and His tags. The untagged RHDV replicon acted as a negative control. After two-step affinity purification, the eluted protein complexes were resolved by sodium dodecyl sulfate polyacrylamide gel electrophoresis (SDS-PAGE) and the protein bands were visualized with silver staining (Fig. 2A). In total, 11 specific or enriched bands were sliced from the RHDV-luc-His_25_/HA_442_ lane and the proteins they contained were identified using mass spectrometry (MS) (Table 1). The identified host proteins were associated with cytoskeleton components, intracellular transport, chaperone, RNP components, and translation machine-related proteins. Among these proteins, numerous proteins have been shown to interact with some single-stranded positive-strand RNA viral proteins to regulate viral replication, such as HnRNPK, HSPA8, DDX5, ANXA2, and PI4KA [19-23] (Fig. 2B and Table 1). In addition, the protein interaction network of the identified components of the RC was mapped using STRING online software (https://string-db.org/). As showed in Fig. 2C, there are intricate interaction networks for the components of the RHDV RC. We found that NCL not only binds to the RHDV replicase RdRp but it also interacts with many host proteins, such as casein kinase II subunit alpha (CSNK2A1), heterogeneous nuclear ribonucleoprotein K (HnRNPK), 40S ribosomal protein S5 (RPS5), 60S ribosomal protein L11 (RPL11), and so on.

**Table 1.** Categories of host factors found to be associated with RHDV replicase^a^

| Category and band no. | Protein score | Mass (kDa) | Gene name | Protein description | No. of unique peptides | No. of peptides | SC (%) |
| --- | --- | --- | --- | --- | --- | --- | --- |
| **RHDV protein** |  |  |  |  |  |  |  |
| 6 | 273.3 | 57.8 | *RdRp* | RHDV RdRp | 3 | 4 | 21.1 |
| 10 | 186.4 | 25.1 | *p23* | RHDV p23 | 2 | 3 | 15.5 |
| 11 | 165.8 | 16.2 | *p16* | RHDV p16 | 2 | 4 | 10.4 |
| **Transport** |  |  |  |  |  |  |  |
| 8 | 57.1 | 39.2 | *ANXA2* | Annexin A2 | 4 | 7 | 10.4 |
| 1 | 201.3 | 233.6 | *PI4KA* | Phosphatidylinositol 4-kinase alpha | 6 | 6 | 30.5 |
| 5 | 303.5 | 68.9 | *ALB* | Serum albumin | 8 | 10 | 15.2 |
| 11 | 287.6 | 15.6 | *HBA* | hemoglobin subunit alpha | 5 | 5 | 40.4 |
| 6 | 173.2 | 59.7 | *ATP5A1* | F-type H+-transporting ATPase subunit beta | 3 | 3 | 7.8 |
| 11 | 365.6 | 16.1 | *HBB* | hemoglobin subunit beta | 7 | 9 | 58.7 |
| 9 | 98.3 | 32.9 | *SLC25A5* | solute carrier family 25 | 3 | 3 | 11.2 |
| 11 | 99.8 | 12.6 | *FABP5* | fatty acid-binding protein 5 | 3 | 4 | 13.5 |
| **Cytoskeleton** |  |  |  |  |  |  |  |
| 11 | 105.5 | 14.3 | *CFL1* | Cofilin 1 | 2 | 6 | 16.8 |
| 3 | 316.5 | 102.9 | *ACTN1* | Actinin alpha 1 | 2 | 7 | 32.4 |
| 4 | 286.3 | 87.4 | *ACTN4* | Actinin alpha 4 | 2 | 6 | 28.5 |
| 8 | 357.8 | 41.7 | *ACTB* | Actin, cytoplasmic 1 | 2 | 5 | 19.0 |
| 7 | 419.4 | 53.6 | *VIM* | Vimentin | 13 | 13 | 25.3 |
| **RNP complex** |  |  |  |  |  |  |  |
| 10 | 78.5 | 25.7 | *HNRNPA1* | Heterogeneous nuclear ribonucleoprotein A1 | 3 | 7 | 13.6 |
| 10 | 67.9 | 30.3 | *HNRNPAB* | Heterogeneous nuclear ribonucleoprotein A/B | 2 | 6 | 8.9 |
| 7 | 127.2 | 50.9 | *HNRNPK* | Heterogeneous nuclear ribonucleoprotein K | 4 | 9 | 11.6 |
| 5 | 87.0 | 67.5 | *DDX5* | DEAD-box helicase 5 | 2 | 4 | 8.3 |
| 5 | 90.6 | 69.4 | *NCL* | Nucleolin | 8 | 10 | 28.8 |
| 2 | 117.8 | 130.7 | *SORBS2* | Sorbin and SH3 domain containing 2 | 5 | 5 | 14.7 |
| 10 | 135.6 | 26.2 | *NUDT21* | cleavage and polyadenylation specificity factor subunit 5 | 3 | 3 | 10.4 |
| 5 | 99.1 | 70.9 | *PABPC1* | Polyadenylate-binding protein 1 | 4 | 4 | 16.3 |
| 9 | 128.4 | 36.0 | *YBX1* | Nuclease sensitive element-binding protein 1 | 5 | 6 | 22.1 |
| 6 | 112.1 | 57.4 | *PTBP1* | Polypyrimidine tract-binding protein 1 | 4 | 7 | 6.7 |
| 6 | 65.2 | 52.1 | *SRSF1* | splicing factor, arginine/serine-rich 1 | 6 | 7 | 8.8 |
| 5 | 75.9 | 63.5 | *CPSF6* | cleavage and polyadenylation specificity factor subunit 6/7 | 3 | 10 | 10.9 |
| 6 | 75.2 | 62.5 | *DDX41* | Probable ATP-dependent RNA helicase | 3 | 12 | 14.5 |
| **Chaperon** |  |  |  |  |  |  |  |
| 7 | 65.2 | 45.1 | *CSNK2A1* | Casein kinase II subunit alpha | 5 | 5 | 7.5 |
| 4 | 236.2 | 71.0 | *HSPA8* | Heat shock cognate 71 kDa protein | 7 | 7 | 14.2 |
| 9 | 153.9 | 30.7 | *PYCR2* | Pyrroline-5-carboxylate reductase | 4 | 5 | 16.8 |
| 3 | 70.1 | 116.8 | *DSG1* | Desmoglein 1 | 2 | 5 | 9.0 |
| 11 | 50.3 | 14.7 | *LYZ* | Lysozyme C | 2 | 3 | 11.5 |
| 4 | 53.5 | 73.5 | *HSPA9* | heat shock protein family A (Hsp70) member 9 | 2 | 2 | 13.8 |
| 10 | 152.2 | 26.2 | *NUDT21* | Nudix hydrolase 21 | 3 | 3 | 24.5 |
| 8 | 142.8 | 40.3 | *PDCD2L* | Programmed cell death protein 2-like | 4 | 4 | 28.7 |
| 11 | 108.3 | 16.0 | *FAM207A* | family with sequence similarity 207 member A | 5 | 8 | 19.6 |

**Table1** (Continued)
| Category and band no. | Protein score | Mass (kDa) | Gene name | Protein description | No. of unique peptides | No. of peptides | SC (%) |
| --- | --- | --- | --- | --- | --- | --- | --- |
| Translation machines |  |  |  |  |  |  |  |
| 7 | 94.2 | 50.0 | *EEF1A* | Elongation factor 1-alpha | 7 | 7 | 9.3 |
| 11 | 135.4 | 17.7 | *RPS18* | Ribosomal protein S18 | 4 | 4 | 18.7 |
| 11 | 113.3 | 16.3 | *AAES* | 40S ribosomal protein S14 | 3 | 3 | 28.3 |
| 10 | 174.3 | 20.2 | *RPL11* | Ribosomal protein L11 | 4 | 4 | 20.1 |
| 10 | 283.1 | 21.2 | *RPL12* | Ribosomal protein L12 | 5 | 5 | 33.5 |
| 11 | 164.9 | 12.8 | *RPL30* | 60S ribosomal protein L30 | 4 | 4 | 50.5 |
| 11 | 68.3 | 15.5 | *RPS17* | 40S ribosomal protein S17 | 3 | 3 | 29.6 |
| 10 | 323.2 | 22.9 | *RPS5* | Ribosomal protein S5 | 7 | 7 | 18.5 |
| 9 | 258.2 | 31.1 | *RPS3* | 40S ribosomal protein S3 | 9 | 9 | 25.8 |
| 7 | 74.4 | 45.3 | *EIF4A1* | Eukaryotic initiation factor 4A-I | 4 | 6 | 9.7 |
| 11 | 122.6 | 13.7 | *RPS25* | Ribosomal protein S25 | 5 | 5 | 21.3 |
| 11 | 75.3 | 14.0 | *RPS15A* | Ribosomal protein S15a | 2 | 5 | 18.3 |

<sup>a</sup> Protein lists the gene name for each of the proteins identified in Fig.2B. SC refers to the percent sequence coverage for the protein.

**Figure 2.**
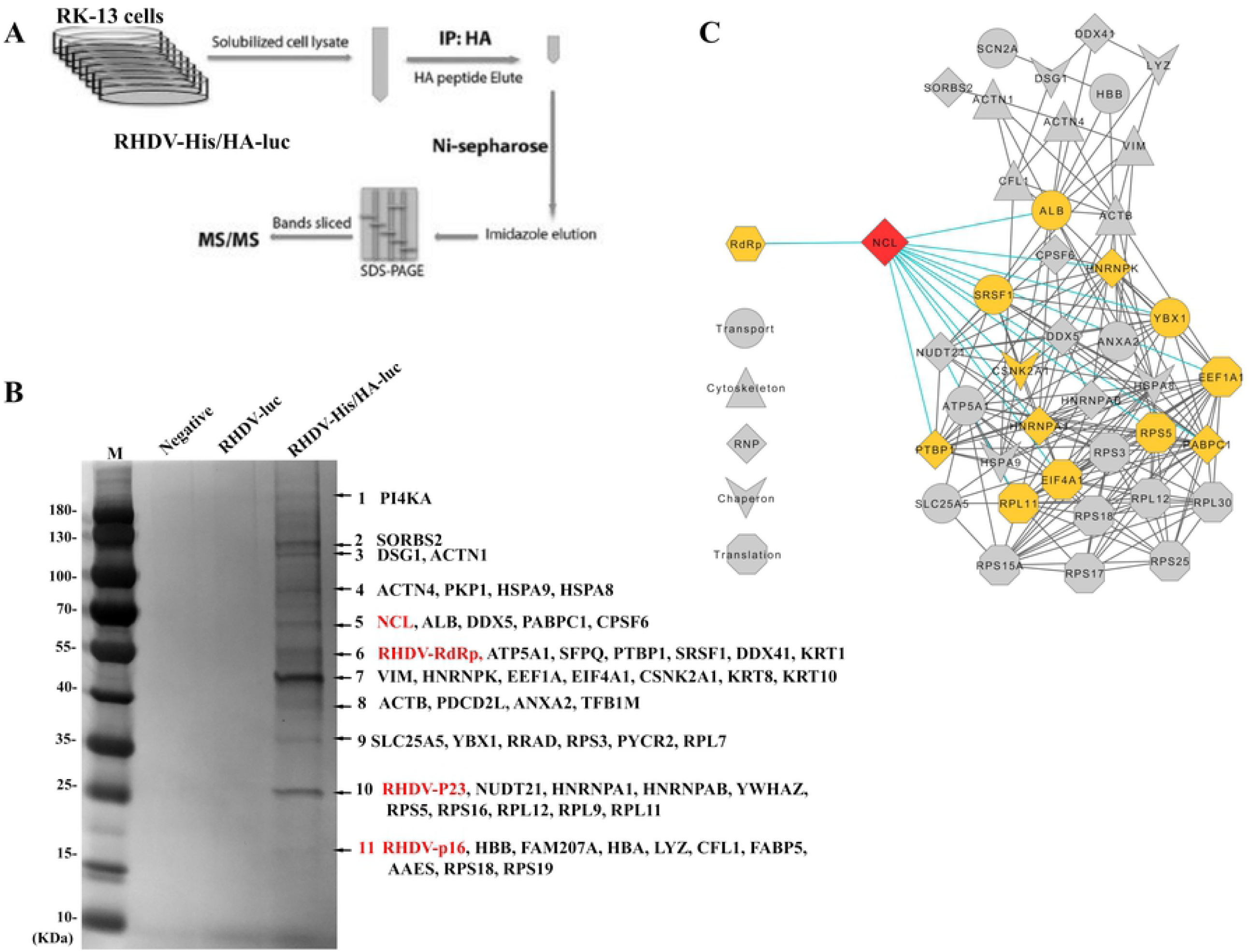
Affinity purification of solubilized RHDV replicase RdRp-associated host factors. **(A)** Schematic of two-step affinity purification of the RHDV replicase RdRp-associated host proteins. **(B)** After two-step affinity purification, the eluted proteins were resolved by SDS-PAGE. Specifically, enriched protein bands (arrows) in the RHDV-His/HA-luc sample were identified by mass spectrometry. Mainly, host proteins and identified viral proteins in the bands indicated are shown. Host factors selected for further study are marked in red. **(C)** Network analysis of RHDV replicase-associated protein. The interaction network shows the proteins identified as being associated with RHDV replicase by affinity purification. To simplify these interdependencies, host factors that did not interact with other factors in the network are not shown. NCL is marked in red. Proteins that interact with NCL are labeled in yellow. Note: the network was generated using the data presented in Table 1. The interaction network was generated using the STRING online tool and then presented using Cytoscape software [67].

### NCL is involved in RHDV replication

NCL is a phosphoprotein that is ubiquitously and abundantly expressed in many eukaryotic cells and highly conserved during evolution, as it is involved in a remarkably large number of cellular activities [24]. In general, NCL is mainly distributed in the nucleolus, but it also exists in the nucleoplasm, cytoplasm, and cell surface, where its specific functions vary, such as ribosome biogenesis, proliferation, and cell cycle regulation. NCL also plays important roles in the replication and intracellular trafficking of multiple viruses [25-31].

To determine if NCL is required for RHDV replication, RK-13 cells were co-transfected with an RHDV replicon, NCL siRNA or Flag-tagged NCL plasmids and internal control plasmid encoding an *Rluc* gene. First, the effect of NCL siRNA on the viability of RK-13 cells was detected using a CCK-8 kit, according to the manufacturer’s instruction. We found that NCL knockdown would have some effect on cell viability and proliferation (Fig. S2A), meanwhile, resulting the expression level of Rluc was affected (Fig. S2B). Since the toxicity, cell number, and transfection efficiency of each treatment group have the same effect on the RHDV replicon and Rluc plasmid, we used Rluc plasmids to eliminate the non-specific effects of siRNA and other treatments on cells, and to correct the experimental data. The reporter luciferase activity was evaluated using a dual-luciferase reporter assay system with cell lysates that were harvested at 24 h and 48 h post-transfection (hpt). Fluc activity was normalized with respect to a co-transfected plasmid encoding an Rluc. Similar results were obtained in three independent experiments. The results showed that there is a positive correlation between the expression level of Fluc and NCL. This decreased with increasing NCL siRNA transfection dose and increased with increased dose of Flag-NCL transfection (Fig. 3A and 3B).

**Figure 3.**
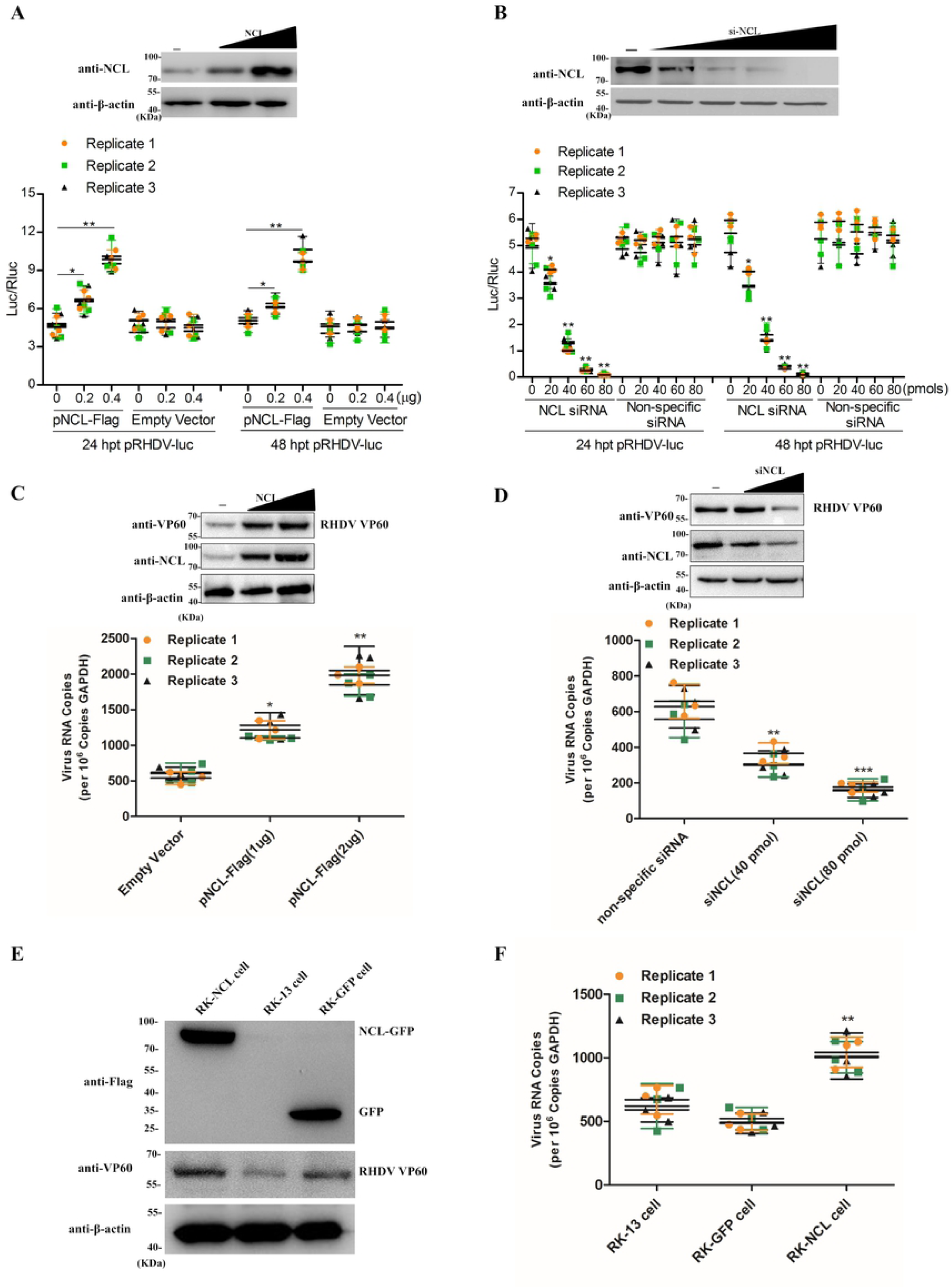
NCL is involved in RHDV replication. **(A)** The effect of NCL eukaryotic plasmids on viral replicon activity. Relative luciferase activity was evaluated in RK-13 cells carrying pRHDV-luc, and trans-supplemented NCL eukaryotic plasmids pFlag-NCL (0.2 μg, 0.4 μg) at 24 hpt and 48 hpt. The p3 × Flag-CMV-14 vector acted as negative control. The luciferase activity in RK-13 cells was evaluated by measuring Fluc activity. Rluc activity was measured to normalize the transfection efficiency. **(B)** The effect of NCL siRNA on viral replicon activity. The RK-13 cells, co-transfected with pRHDV-luc and NCL siRNA (20 pmol, 40 pmol, 60 pmol, or 80 pmol), were lysed at 24 hpt and 48 hpt, and Fluc activity was measured based on RLUs and normalized according to the results obtained for a co-transfected pLTK plasmid encoding Rluc. The nonspecific siRNA acted as negative control. **(C-D)** The effect of NCL on mRHDV replication. The RK-13 cells, transfected with pFlag-NCL(1μg, 2μg) or NCL siRNA (40 pmol, 80 pmol), were infected with mRHDV (MOI = 1) at 24 hpt, and the level of mRHDV replication were evaluated by WB and qRT-PCR at 48 hpi. The p3 × Flag-CMV-14 vector and nonspecific siRNA acted as negative control. **(E-F)** The replication ability of mRHDV in RK-NCL cells. The expression level of NCL in RK-NCL cells at 10 passages was determined by western blot analysis with anti-Flag mAb. The RHDV replication levels in RK-NCL cells infected with mRHDV (MOI = 1) were evaluated by WB and qRT-PCR at 48 hpi. RK-GFP cells acted as negative controls; RK-13 cells acted as blank controls. Student *t*-tests and analysis of variance were used for statistical analyses. \**p* < 0.05 and \*\**p* < 0.01. The number of cells used in all replicate experiments was similar.

Subsequently, we examined the effect of NCL on mRHDV, which could proliferate in RK-13 cells [15]. We also used NCL siRNA or Flag-tagged NCL to change the expression level of NCL, and then infected with mRHDV (MOI = 1). At 48 hpi, the replication level of mRHDV were detected by WB and qPCR. The results were similar to RHDV replicons. As shown in Fig. 3C and 3D, the replication level of mRHDV increased with increased dose of Flag-NCL and decreased with increasing NCL siRNA.

In addition, we successfully constructed an RK-NCL cell line, which overexpressed the *NCL* gene, using a lentiviral packaging system (Fig. 3E). To evaluate the replication dynamics of mRHDV in RK-NCL cells, the cells were infected with mRHDV (MOI=1), and subsequently the expression level of VP60 was evaluated with qRT-PCR and WB at 48 hpi. The results showed that the expression level of VP60 in RK-NCL cells was significantly higher than that in control cells (RK-GFP cells and RK-13 cells) (Fig. 3E and 3F). Collectively, these data suggest that NCL is involved in RHDV replication.

### NCL interaction with RHDV RdRp

NCL is a phosphoprotein with protein-binding activity, and it has been previously reported that NCL regulates viral replication by binding to viral proteins [32-40]. To determine if NCL regulates RHDV replication through interaction with viral nonstructural proteins, we used mammalian two-hybrid (M2H) assays to screen the interaction between NCL and viral nonstructural proteins. As shown in Fig. 4A, NCL interacted with RdRp, p16, and p23. To determine whether endogenous NCL binds to these viral nonstructural proteins, during RHDV genome replication, we assessed the interaction between NCL and these viral proteins in RK-13 cells, in the presence and absence of mRHDV infection for 24 h at 37 °C. The results of an immunoprecipitation (IP) assay performed with cell lysates using NCL mAb showed that regardless of whether the cell lysates were treated with RNase, NCL interacted with RdRp, p16, and p23 in infected cells, but did not in uninfected cells (Fig. 4B).

**Figure 4.**
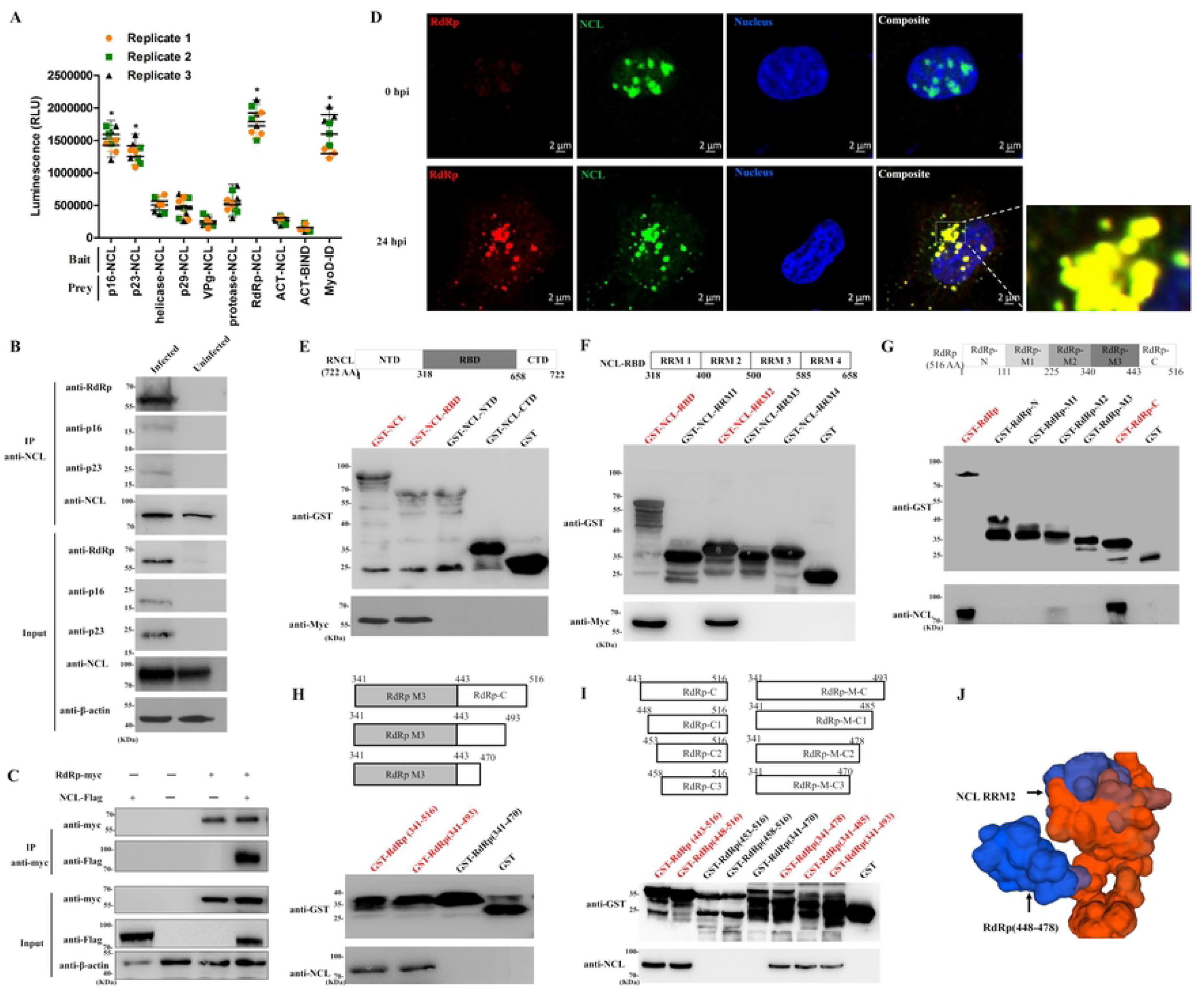
NCL interacts with RHDV RdRp. **(A)** M2H interaction of NCL with RHDV nonstructural proteins. Student *t*-tests and analysis of variance were used for statistical analyses. \**p* < 0.05 and \*\**p* < 0.01. The number of cells used in all replicate experiments was similar. **(B)** NCL binds to RdRp, p16 and p23 during RHDV replication. An IP assay was performed on cell lysates using NCL mAb in RK-13 cells that were infected or uninfected with mRHDV, then immunoblotted with Abs against NCL, RdRp, p16 or p23. β-actin was used as an internal control. Cells uninfected with mRHDV served as negative controls. **(C)** Validation of the interaction of RHDV RdRp with NCL in a Co-IP assay. RK-13 cells were co-transfected with the indicated plasmids (+) or empty vectors (-). At 48 hpt, cells were lysed, and IP of myc-fused proteins was performed using anti-myc mAb. Lysates (input) and IPs were analyzed with IB using antibody against myc or Flag. β-actin was used as an internal control. **(D)** Confocal microscopy analysis of NCL (green) and RdRp (red) in RK-13 cells infected with mRHDV at 24 hpi with mAbs against NCL and RdRp. The small white boxes represent amplified random co-localization spots within the merged image, and the co-localization spots are indicated with white arrowheads. **(E–F)** The functional domain of NCL interacting with RdRp was identified by GST pull-down assay. GST fusions of various NCL domains were used as bait; RdRp expressed in RK-13 cells, obtained at 48 hpt with pRdRp-myc plasmids, was used as prey. RdRp binding was immunoblotted with anti-myc mAb. The GST protein acted as a negative control. **(G–I)** IB analysis of the glutathione affinity pull-down assays was performed to map the binding domain of the RdRp protein. We used GST-tagged RdRp domains as bait and NCL expressed in RK-13 cells as prey. After extensive washing, NCL binding was determined by IB with anti-Flag mAb. The GST protein acted as a negative control. The interactions are shown in red. (**J**) Modeling of the functional area of NCL-RdRp interaction.

RdRp is a replicase of RHDV that plays a key role in viral replication [41]. To prove NCL interacts with RdRp, co-immunoprecipitation (Co-IP) assays were used with a myc mAb in RK-13 cells, which were co-transfected with pRdRp-myc and pNCL-Flag eukaryotic expression plasmids. We showed that overexpressed NCL-Flag was present in the anti-myc (RdRp-myc) immunocomplex (Fig. 4C). Furthermore, NCL interacted with RdRp in RHDV-infected cells (Fig. 4B). These results showed that NCL can interact with RHDV RdRp. Moreover, an immunofluorescence assay (IFA) was performed using mAbs against NCL and RdRp in RK-13 cells infected with mRHDV at 24 hours post infection (hpi). As shown in Fig. 4D, NCL was co-localized with RHDV RdRp in the RK-13 cell cytoplasm. In addition, the distribution of NCL in the cytoplasm increased after RHDV infection.

The multifunctionality of NCL mainly results from its multidomain structure, which is composed of three main structural domains: the N-terminal domain (NTD), the central domain, and the C-terminal domain (CTD). The central domain contains fours RRMs, which are also called RNA-binding domains (RBDs). To identify the functional domain of NCL for NCL-RHDV RdRp interactions, GST fusion proteins corresponding to NCL and subfragments (GST-NCL-NTD, GST-NCL-RBD, and GST-NCL-CTD, respectively) were prepared for use as bait proteins in GST pull-down assays, to determine their abilities to interact with the RdRp protein expressed in RK-13 cells. As shown in Fig. 4E, GST-NCL and GST-NCL-RBD bound to RdRp whereas the other proteins were undetectable. This result confirmed that binding to RdRp requires the RBD domain of NCL. To further map the NCL RBD motif responsible for NCL-RdRp interactions, we prepared GST fusion proteins corresponding to subfragments of the NCL-RBD domain, including NCL-RRM1, NCL-RRM2, NCL-RRM3, and NCL-RRM4. A set of pull-down assays showed that GST-NCL-RBD and GST-NCL-RRM2 interact with RdRp whereas the other proteins did not (Fig. 4F). These findings indicate that NCL interacts with RHDV RdRp via the RRM2 motif.

To characterize the critical domain of RdRp for NCL-RdRp interactions, the GST fusion proteins corresponding to RdRp and subfragments of RdRp (GST-RdRp, GST-RdRp-N, GST-RdRp-M1, GST-RdRp-M2, GST-RdRp-M3, and GST-RdRp-C) were prepared for use as bait proteins in GST pull-down assays, to determine their ability to interact with the NCL protein expressed in RK-13 cells. The results of these assays showed that GST-RdRp, GST-RdRp-C and GST-RdRp-M2 bound to NCL, but the other proteins did not, and the binding ability of GST-RdRp-M2 to NCL is very weak (Fig. 4G). Subsequently, we prepared subfragments of RdRp_(341-516)_ GST fusion proteins, including GST-RdRp_(341-493)_, and GST-RdRp_(341-470)_ for use as bait proteins in GST pull-down assays. As shown in Fig. 4H, GST-RdRp_(341-516)_ and GST-RdRp_(341-493)_ bound to NCL whereas GST-RdRp_(341-470)_ did not. In addition, RdRp_(443-516)_ was split into the fragments RdRp_(448-516)_, RdRp_(453-516)_, and RdRp_(458-516)_; and RdRp_(341-493)_ was split into the fragments RdRp_(341-485)_, RdRp_(341-478)_, and RdRp_(341-470)_, which were fused with GST and expressed. The results of further pull-down assays showed that GST-RdRp_(443-516)_, GST-RdRp_(448-516)_, GST-RdRp_(341-493)_, GST-RdRp_(341-485)_, and GST-RdRp_(341-478)_ bound to NCL whereas the other proteins did not (Fig. 4I). These results suggest that NCL directly and specifically interacts with the C-terminal aa residues 448–478 of RHDV RdRp.

In addition, as predicated using the SWISS-MODEL online tool, we found that there is a “key” and “lock” structure formed by the amino acid sequence 448-478 of RdRp and RRM2 of NCL, providing space for the interaction between NCL and RdRp (Fig. 4J). Moreover, analysis of the RdRp sequence of GI.1a–GI.1d genotypes of RHDV showed that the C-terminal aa residues 448–478 were highly conserved (Fig. S3A). Together, these observations confirm that RHDV RdRp interacts with the RRM2 motif of NCL via the C-terminal aa residues 448–478, which is a conserved sequence in RHDV.

### NCL interaction with RHDV p16

The function of p16 in RHDV remains unclear. Previous studies have reported that p16 can accumulate in subnuclear compartments, which may point to a specific interaction with nucleic acids and/or cellular proteins [42].

NCL is one of the most abundant proteins in the nucleolus. To investigate whether NCL directly interacts with p16, a Co-IP assay was performed with a myc mAb in RK-13 cells, which were co-transfected with p16-myc and NCL-Flag eukaryotic expression plasmids. IB analysis using a mAb against the Flag tag showed a band corresponding to NCL in the myc Co-IP assay, indicating a direct interaction between RHDV p16 and NCL (Fig. 5A). Moreover, we conducted an IFA using NCL mAbs and p16 polyclonal antibody in RK-13 cells infected with mRHDV at 24 hpi. As shown in Fig. 5B, NCL was co-localized with RHDV p16 in the RK-13 cell nucleolus and cytoplasm.

**Figure 5.**
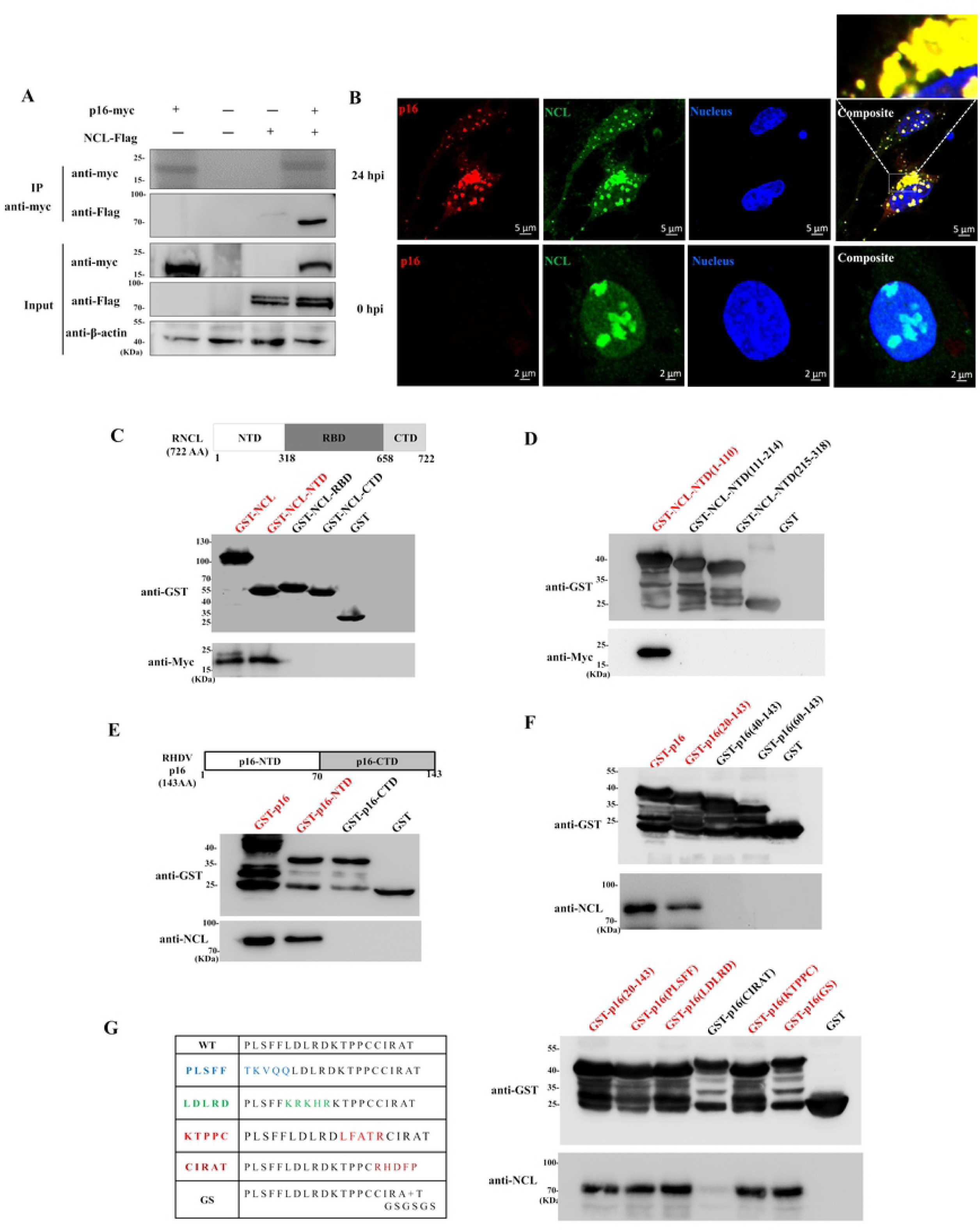
NCL interacts with RHDV p16. **(A)** We identified the interaction between RHDV p16 and NCL using Co-IP assay. RK-13 cells were co-transfected with the indicated plasmids (+) or empty vectors (-). Cell lysates were prepared 48 h post-transfection and the proteins were subjected to IP followed by IB analysis. **(B)** Confocal microscopy analysis of NCL (green) and RdRp (red) in RK-13 cells infected with mRHDV at 24 hpi with Abs against NCL and p16. The small white boxes represent amplified random co-localization spots within the merged image, and the co-localization spots are indicated with white arrowheads. **(C–D)** The functional domain of NCL interacting with p16 was identified by GST pull-down assays. GST fusions of various NCL domains were used as bait, and myc fusion-p16 proteins expressed in RK-13 cells, were used as prey. p16 binding was immunoblotted with anti-myc mAb. The GST protein acted as a negative control. **(E–G)** Glutathione affinity pull-down assays were performed to map the binding domain of p16 protein. We used GST-tagged p16 domains as bait and Flag fusion-NCL proteins expressed in RK-13 cells as prey. After extensive washing, NCL binding was determined by IB with anti-Flag mAb. The GST protein acted as a negative control. The interactions are shown in red.

To investigate the functional areas of NCL in NCL-RHDV p16 interaction, the GST-NCL fusion protein and its subfragments were used as bait proteins in GST pull-down assays to determine their ability to interact with the p16 protein. As shown in Fig. 5C, only GST-NCL and GST-NCL-NTD bound p16. This result confirms that binding to p16 requires the NTD domain of NCL. Subsequently, we prepared subfragments of NCL-NTD GST fusion proteins, including NCL-NTD_(1-110)_, NCL-NTD_(111-214)_, and NCL-NTD_(215-318)_ for use as bait proteins in a set of GST pull-down assays. As shown in Fig. 5D, GST-NCL-NTD and GST-NCL-NTD_(1-110)_ interact with p16 whereas the other proteins did not. Our data indicate that NCL interacts with RHDV p16 via N-terminal residues 1–110.

Next, we identified the critical domain of p16 for NCL-p16 interactions using a series of glutathione pull-down assays. First, the GST fusion proteins corresponding to p16 and subfragments of p16 (GST-p16, GST-p16_(1-70)_, GST-p16_(20-143)_, GST-p16_(40-143)_, GST-p16_(60-143)_, and GST-p16_(70-143)_) were prepared for use as bait proteins in GST pull-down assays, to determine their ability to interact with NCL protein expressed in RK-13 cells. The results of these assays showed that GST-p16, GST-p16_(1-70)_ and GST-p16_(20-143)_ bound to NCL, but the other proteins did not (Fig. 5E-F). Subsequently, to pinpoint the key aa responsible for binding of RHDV p16 with NCL, blocks of five aa substitutions were introduced within and beyond the conserved sequence motif. The following non-conservative substitutions to the GST-tagged RHDV p16 protein were made: ^20^PLSFF^24^TKVQQ, ^25^LDLRD^29^KRKHR, ^30^KTPPC^34^LFATR, and ^35^CIRAT^39^RHDFP. Furthermore, GSGSGS was inserted after aa residue 38. It has been reported that GSGSGS is a flexible peptide used in different expression systems to separate functional proteins [43]. Wild-type and mutant RHDV p16 proteins were used as bait proteins in the GST pull-down assays to determine their ability to bind NCL. As shown in Fig. 5G, the ^35^CIRAT^39^RHDFP mutation was not able to bind NCL; the binding capacities of the other mutations to the residues at positions 20–40 were partially reduced. In addition, analysis of the p16 sequence of the GI.1a–GI.1d genotypes of RHDV showed that the ^35^CIRAT^39^ motif was highly conserved (Fig. S3B). These observations confirm that NCL interacts with RHDV p16 via aa residues 1–110 of the NTD domain of NCL bound to the ^35^CIRAT^39^ motif of p16.

### NCL interaction with RHDV p23

The function of p23 of RHDV is also unclear. Previous studies have reported that RHDV p23 is similar to other caliciviruses, which show an endoplasmic reticulum-like localization pattern [42]. It is well known that the murine norovirus (MNV), human norovirus (NV), and feline calicivirus (FCV) homologues of p23 play a role in the induction of intracellular membrane rearrangements associated with viral replication [44-46]. As previously described, NCL may have a direct role in the assembly of the ribosomal subunits by bringing together ribosomal proteins and RNA [47].

To identify whether p23 directly interacts with NCL, a Co-IP assay was used with a myc mAb in RK-13 cells, which were co-transfected with p23-myc and NCL-Flag eukaryotic expression plasmids. IB analysis using a mAb against the Flag tag showed a band corresponding to NCL in the myc Co-IP assay, indicating a direct interaction between RHDV p23 and NCL (Fig. 6A). In addition, we performed an IFA using mAbs against NCL and polyclonal antibody against p23 in RK-13 cells infected with mRHDV at 24 hpi. The result showed that NCL was co-localized with p23 in the RK-13 cell cytoplasm (Fig. 6B).

**Figure 6.**
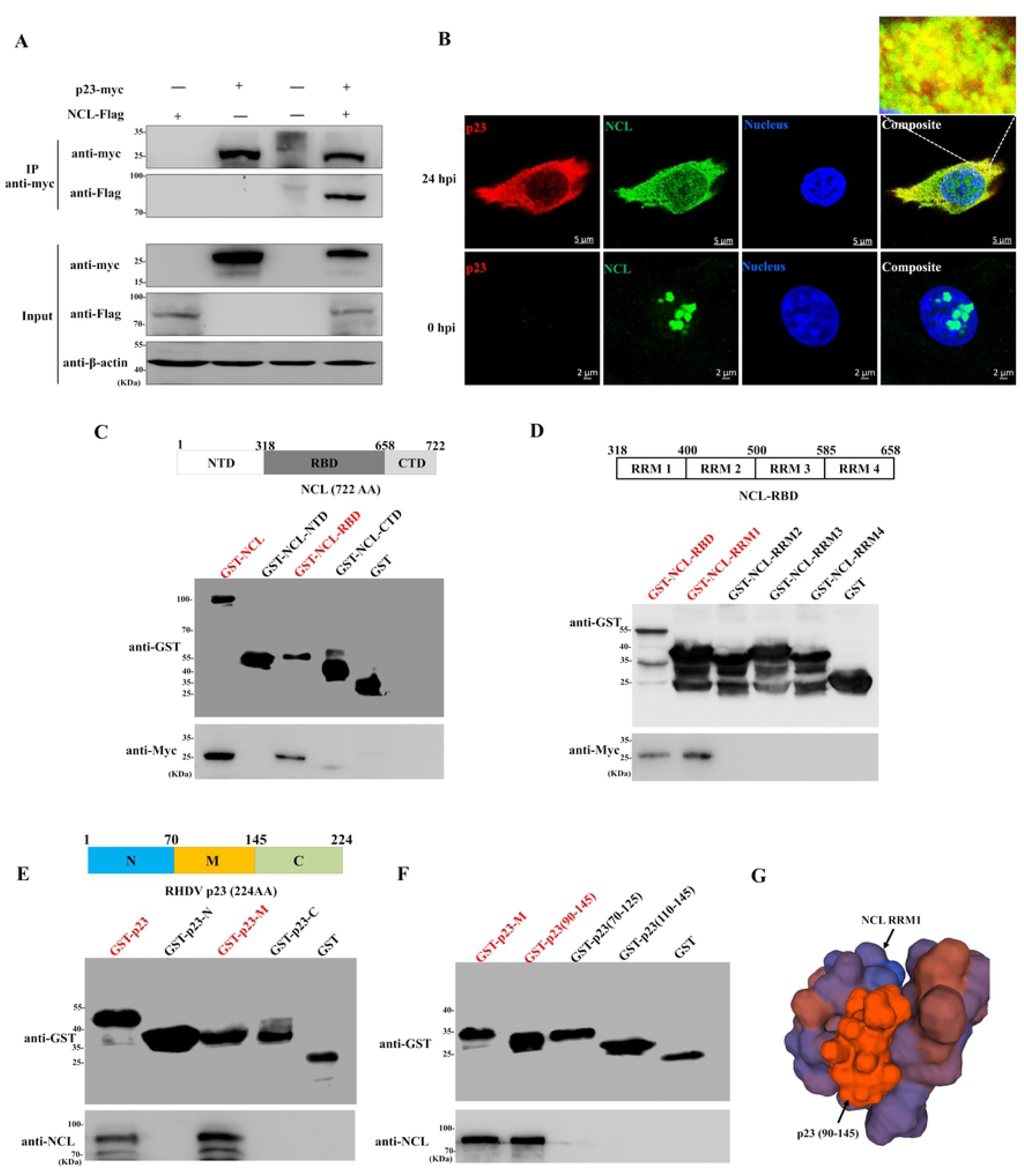
NCL interacts with RHDV p23. **(A)** Co-IP of RHDV p23 with NCL in RK-13 cells. RK-13 cells were transfected with the indicated plasmids (+) (myc-p23 or Flag-NCL) and empty vectors (-) for 48 h. Cell lysates were incubated with anti-myc mAb-coated beads and Co-IP proteins were subjected to IB analysis. **(B)** Confocal microscopy analysis of NCL (green) and p23 (red) in RK-13 cells infected with mRHDV at 24 hpi with Abs against NCL and p23. The small white box represents amplified random co-localization spots within the merged image, and the co-localization spot is indicated with a white arrowhead. **(C–D)** The functional domain of NCL interacting with p23 was identified using GST pull-down assays. GST fusions of various NCL domains were used as bait, and myc fusion-p23 proteins expressed in RK-13 cells were used as prey. p23 binding was immunoblotted with anti-myc mAb. The GST protein acted as a negative control. **(E–F)** Glutathione affinity pull-down assays were performed to map the binding domain of p23 protein. We used GST-tagged p23 domains as bait and Flag fusion-NCL proteins expressed in RK-13 cells as prey. After extensive washing, NCL binding was determined by IB with anti-Flag mAb. The GST protein acted as a negative control. The interactions are shown in red. (**G**) Modeling of the functional area of NCL-p23 interaction.

To characterize the critical domain of NCL for NCL-RHDV p23 interactions, the GST fusion proteins corresponding to NCL and subfragments were used as bait proteins in GST pull-down assays to determine their ability to interact with p23 protein expressed in RK-13 cells. As shown in Fig. 6C, both GST-NCL and GST-NCL-RBD bound to RHDV p23, but the other proteins did not. Subsequently, the subfragments of NCL-RBD GST fusion proteins were used as bait proteins in a GST pull-down assay. As shown in Fig. 6D, GST-NCL-RRM1 and GST-NCL-RBD bound to RHDV p23 whereas the other proteins did not. Together, these results suggested that RHDV p23 directly and specifically interacts with RRM1 motif of NCL.

To map the RHDV p23 segments responsible for p23-NCL interactions, GST fusion proteins corresponding to subfragments of p23 (p23_(1-70)_, p23_(70-145)_, and p23_(145-224)_) were prepared for use as bait proteins in GST pull-down assays to determine their ability to interact with NCL protein expressed in RK-13 cells. Results of a set of pull-down assays showed that GST-p23_(70-145)_ and GST-p23 interact with NCL (Fig. 6E). Subsequently, subfragments of p23_(70-145)_ GST fusion proteins including GST-p23_(90-145)_, GST-p23_(110-145)_, and GST-p23_(70-125)_ were prepared for use as bait proteins in a GST pull-down assay. As shown in Fig. 6F, GST-p23_(90-145)_ and p23_(70-145)_ bound to NCL whereas the other proteins did not. Our data indicate that RHDV p23 interacts with NCL via aa residues 90–145. Moreover, as predicated using the SWISS-MODEL online tool, we found that there is a “mortise” and “tenon” structure formed by the amino acid sequence 90-145 of p23 and RRM1 of NCL, which providing a structural basis for the interaction between NCL and p23 (Fig. 6G). In addition, analysis of the p23 sequence of GI.1a–GI.1d genotypes of RHDV showed that the aa sequence 90–145 of RHDV p23 is conserved (Fig. S3C). These results suggest that NCL interacts with RHDV p23 via the RRM1 motif of NCL bound to the aa sequence 90–145 of p23.

### NCL is required for RHDV replication

The above results fully demonstrated that NCL binds to the RdRp, p16, and p23 proteins of RHDV. To explore the role of these interactions in RHDV replication, a series of recombinant RHDV replicons were obtained by mutating or replacing the region that interacts with NCL in the wild-type RHDV replicon. The luciferase activity of RK-13 cells transfected with these recombinant RHDV replicons were analyzed and compared. The abundance of Fluc mRNA in the absence or presence of these binding sites, interacting with NCL, was evaluated by qRT-PCR. Our results showed that the expression levels of Fluc derived from RHDV-luc/ΔRdRp, RHDV-luc/ΔRdRp_(448-478)_, RHDV-luc/Δp16, RHDV-luc/Δp16_(35-38)_, RHDV-luc/Δp23, and RHDV-luc/Δp23_(90-145)_ were approximately 7%, 9%, 51%, 52%, 46%, and 44% respectively, compared with RHDV-luc (Fig. 7A). Subsequently, using the lentiviral packaging system, we successfully obtained RK-NCL-NTD, RK-NCL-RBD, and RK-NCL-CTD cell lines stably expressing the NCL NTD, RBD, and CTD domains, respectively (Fig. S4). We found that the replication levels of the RHDV replicon and mRHDV were significantly increased in RK-NCL and RK-NCL-RBD cells and significantly decreased in RK-NCL-NTD cells, but not in RK-NCL-CTD or RK-GFP cells (Fig. 7B and 7C). In addition, to better reflect the effects of NCL domains on RHDV replication, we inoculated mRHDV (MOI = 0.1) at low doses in each cell line. The mRHDV was constructed in our previous study and it has been determined that the virus can efficiently replicate in RK-13 cells [15]. The results showed that the level of replication of mRHDV in each cell line was similar to that of the RHDV replicon. From the above results, we revealed that the RBD domain is a functional domain that binds NCL to RdRp and p23, and the NTD domain is an active region that binds NCL to p16. Therefore, NCL segments that bind to the nonstructural proteins of RHDV play an important role in RHDV replication.

**Figure 7.**
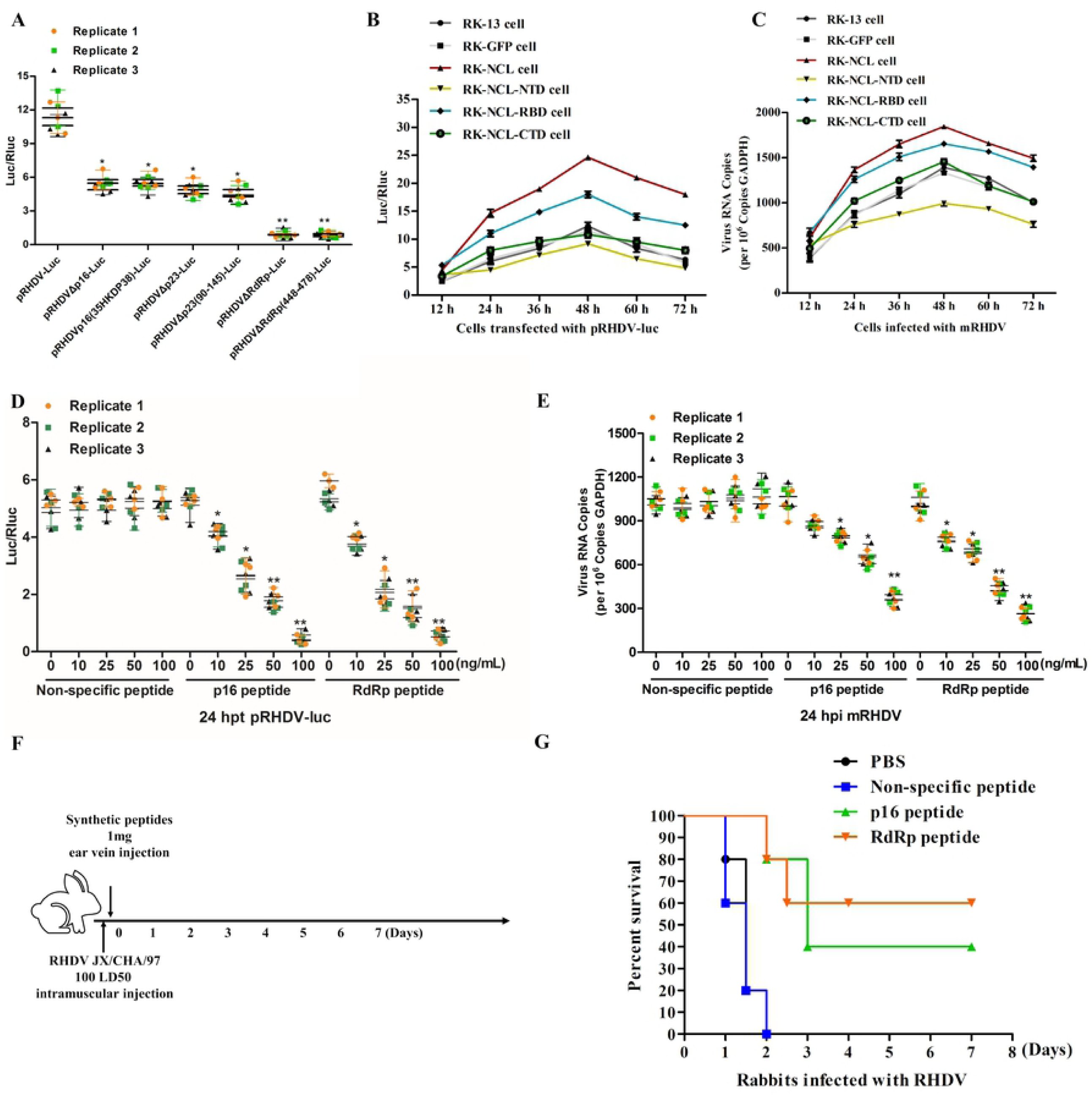
Interactions between NCL and RdRp, p16, and p23 are required for RHDV replication. **(A)** The RHDV replicon and its mutants were transfected with RK-13 cells for 24 h and then lysed to detect luciferase activity in the lysate. Rluc activity was measured to normalize the transfection efficiency. **(B)** Relative luciferase activity in RK-NCL-NTD cells, RK-NCL-RBD cells, and RK-NCL-CTD cells carrying pRHDV-luc at 12, 24, 36, 48, 60, and 72 h post-transfection. RK-NCL cells acted as positive controls; RK-GFP cells acted as negative controls; RK-13 cells acted as blank controls. **(C)** RHDV mRNA levels in RK-NCL cells infected with mRHDV (MOI = 0.1) were evaluated by qRT-PCR at 12, 24, 36, 48, 60, and 72 hpi. RK-GFP cells acted as negative controls; RK-13 cells acted as blank controls. **(D)** Relative luciferase activity was evaluated in RK-13 cells carrying pRHDV-luc, treated with p16 peptide, RdRp peptide, non-specific peptide, or PBS, at 24 hpt. The non-specific peptide and PBS acted as negative controls. **(E)** The effect of RK-13 cell treated with p16 peptide, RdRp peptide, non-specific peptide or PBS on RHDV replication. The mRNA levels in RK-13 cells infected with mRHDV (MOI = 0.1) were evaluated by qRT-PCR at 24 hpi. Student *t*-tests and analysis of variance were used for statistical analyses. \**p* < 0.05 and \*\**p* < 0.01. The number of cells used in all replicate experiments was similar. **(F)** Schematic diagram of animal experiments. **(G)** Survival of rabbits infected with RHDV JX/CHA/97. All rabbits were challenged intramuscularly with 100 LD50 of RHDV. At 2 hpi, rabbits were injected with RdRp peptide (1 mg), p16 peptide (1 mg), non-specific peptide (1 mg), or PBS, via the ear vein, and subsequently clinically examined daily for 7 days.

Moreover, we blocked the binding site of NCL-RdRp or NCL-p16 with a synthesized peptide (RdRp peptide: ERGVQLEELQVAAAAHGQEFFNFVRKELER; p16 peptide: PLSFFLDLRDKTPPCCIRAT, respectively) and examined the effect on the RHDV replicon and mRHDV replication. As shown in Fig. 7D and 7E, the replication level of the RHDV replicon and mRHDV were all drastically reduced in RK-13 cells treated with the RdRp peptides or p16 peptides, and there was a negative correlation with the dose of the synthesized peptide. Of course, the non-specific peptides (HKFGPVCLCNRAYIHDCGRW) had no effect on the replication of RHDV. We also found that these peptides have a greater effect on the replication of RHDV replicon than mRHDV. Although the RHDV replicon is a replication model of the virus, it lacks capsid protein and cannot be assembled into viral particles. Therefore, the RHDV replicon may not fully reflect the replication process of the virus in the cell. We speculate that there are some other mechanisms that regulate RHDV replication.

In addition, the above experimental results were conducted using RK-13 cells infected with mRHDV or transfected with the RHDV replicon. To assess the role of NCL-RdRp and NCL-p16 interactions during infection with wild-type RHDV, rabbits were injected with the RdRp peptide, p16 peptide, non-specific peptide or PBS, immediately after infection with wild-type RHDV. Over the next 7 days, we counted deaths among experimental rabbits (Fig. 7F). As shown in Fig. 7G, the survival rates of rabbits against virulent RHDV in the groups jnjected with RdRp peptide and p16 peptide was 60% and 40%, respectively. However, all rabbits injected with in the non-specific peptide group and PBS group (negative controls) died within 24–48 hpi with virulent RHDV. These negative control animals exhibited clinical symptoms of RHDV infection.

Together, these results suggest that the interactions between NCL and the nonstructural proteins of RHDV (RdRp, p16, and p23) have important roles in RHDV replication. Importantly, NCL is required for RHDV replication.

### NCL is a link in RHDV replication complex (RC) formation

The positive-strand RNA viruses share a conserved replication mechanism in which viral proteins induce host membrane modification to assemble membrane-associated viral RCs [12]. Viruses hijack host factors to facilitate this energy-unfavorable process [13]. Therefore, the components of the viral RC are numerous and complex. NCL is capable of binding to nonstructural proteins (RdRp, p16, and p23) of RHDV and is involved in the formation of RHDV RCs.

To investigate the specific role of NCL in the formation of RHDV RCs, M2H assays were used to screen the interactions between viral nonstructural proteins and multiple host factors in RCs. As shown in Fig. 8A, there are complex interactions between viral nonstructural proteins and host factors in RCs. For example, p16 interacts with itself, helicase, p29, protease, and NCL; p23 binds to protease and NCL; p29 binds to helicase and VPg; helicase interacts with itself; VPg binds to protease; protease interacts with itself and RPS5; RdRp interacts with RPS5; and NCL binds to HnRNPK, CSNK2A1, RPS5, and RPL11. We subsequently used a series of Co-IP assays with a myc mAb in RK-13 cells, which were co-transfected with bait (myc fusion protein) and prey (Flag fusion protein) eukaryotic expression plasmids. IB analysis using a mAb against Flag showed the specific band corresponding to prey proteins in the myc Co-IP assay (Fig. 8B). These results reveal that RHDV replicase RdRp cannot directly bind to other nonstructural proteins of the virus. It is noteworthy that NCL directly interacts with RHDV RdRp and nonstructural proteins (p16 and p23).

**Figure 8.**
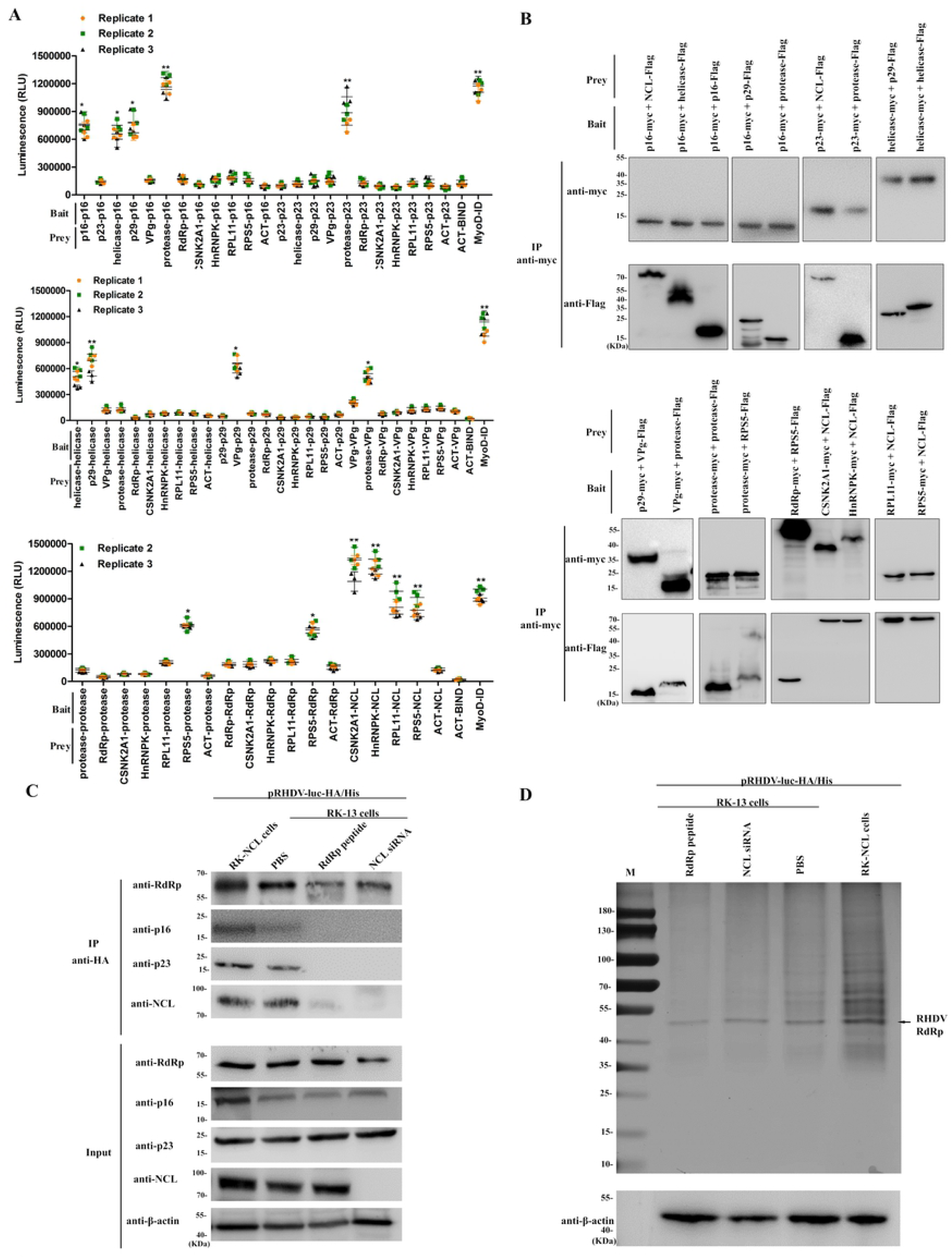
Identification of interactions between RHDV nonstructural proteins and host factors of RCs. **(A)** Identification of these interactions by M2H assays. Bait and prey plasmids were co-transfected with pG5luc plasmids into subconfluent 293T cells at a molar ratio of 1:1:1 for the pACT: pBIND: pG5luc vector. At 48 h after transfection, the 293T cells were lysed, and Rluc and Fluc activities were evaluated using the Promega Dual-Luciferase Reporter Assay System. Student *t*-tests and analysis of variance were used for statistical analyses. \**p* < 0.05 and \*\**p* < 0.01. The number of cells used in all replicate experiments was similar. **(B)** These interactions were verified using Co-IP assays. RK-13 cells were co-transfected with bait and prey plasmids. Cell lysates were prepared 48 h post-transfection and the proteins were subjected to IP followed by IB analysis. myc fusion proteins acted as bait proteins and Flag fusion proteins acted as prey proteins. **(C)** Effect of RdRp peptide and NCL siRNA on p23 and p16 in the RHDV RC. A series of HA tag affinity purification analyses were performed by transfection with the RHDV-luc-His/HA replicon in RK-13 cells, which were treated with RdRp peptide, NCL siRNA or PBS respectively, and in RK-NCL cells. Subsequently, the eluate proteins were subjected to IB analysis using RdRp, p16, p23 or NCL antibodies. PBS acted as a negative control; β-actin acted as an internal control. **(D)** RdRp peptide and NCL siRNA inhibited the formation of the RHDV RC. After HA tag affinity purification, the eluted proteins were resolved by SDS-PAGE. The protein bands were visualized with silver staining. PBS acted as a negative control; β-actin acted as an internal control and was detected by IB with mAb against β-actin.

To test the hypothesis that nucleolin acts as a platform for the RdRp to be attracted to the p16 and p23 proteins, a series of HA tag affinity purification analyses were performed by transfection with the RHDV-luc-His/HA replicon in RK-13 cells that were treated with RdRp peptide, NCL siRNA, or PBS, and in RK-NCL cells. Using IB to detect the purified RdRp-associated protein, we found that the content of purified p16 and p23 in the RdRp peptide- or NCL siRNA-treated cells was significantly reduced or even lost in the RC, and partially increased in RK-NCL cells (Fig. 8C). In addition, the eluted protein complexes were resolved by SDS-PAGE and the protein bands were visualized with silver staining. As shown in Fig. 8D, the RdRp-associated protein content was significantly reduced in RdRp peptide- or NCL siRNA-treated cells and significantly increased in RK-NCL cells.Together, these data suggest that RHDV completes its replication process by hijacking NCL to recruit other viral proteins and host factors, to thus assemble the RHDV RC.

## Discussion

Positive-strand RNA viruses encompass more than one-third of known virus genera and include many medically and practically important human, animal, and plant pathogens [48]. At the outset of infection, positive-strand RNA virus genomes are used as templates for viral protein synthesis in cytoplasm. Subsequently, viral protein recruits host factors to form an RC and redirects the viral genome to function as mRNA, serving as a template for synthesizing new positive-strand gRNA and subgenomic mRNA [48]. Therefore, identification of the components of the viral RC and the interrelationships among the various components, particularly those that interact with viral replicase, will help in understanding the molecular mechanisms of viral replication.

As an important member of the positive-strand RNA viruses, the *Caliciviridae* family has attracted increasing attention because it contains many viruses that infect a wide spectrum of hosts and are a growing threat to human and animal health. However, most caliciviruses cannot be cultured *in vitro*, including some important pathogens such as RHDV and NV; therefore, the replication and pathogenic mechanisms of these viruses remain poorly understood. The emergence and advancement of reverse genetic manipulation technology has provided an excellent operating platform for revealing the molecular mechanism of calicivirus replication. For example, using a NV replicon, a series of works have been carried out to reveal the mechanism of NV replication [49-53]. Recently, we also successfully established an RHDV replicon operating platform [14] and have applied it to study the genomic structure and function of RHDV.

In this study, we purified viral replicase and identified the replicase-associated host factors using an RHDV replicon system in which two different affinity tags were simultaneously inserted in-frame into RdRp. We determined that NCL plays a key role in the formation of RHDV RCs. On the one hand, NCL binds to RHDV replicase (RdRp) (Fig. 4). Similar to other positive-sense RNA viruses, RHDV RdRp has a central role in the viral replication cycle. RdRp has many enzymatic properties; it binds template RNAs, initiates replication, catalyzes elongation, and terminates replication [54]. Moreover, RdRp is able to induce the redistribution of Golgi membranes in kidney and liver cell lines of three different species [55]. Here, our data suggested that interaction between NCL and RdRp is required for RHDV replication (Fig. 7). On the other hand, NCL also interacts with some important host factors, such as HNRNPK, RPL11, CSNK2A1, and RPS5 (Figs. 2, 8, 9). Previous studies have shown that these proteins are involved in the replication of various viruses. For example, HNRNPK has been reported to recognize the 5’ terminal sequence of HCV RNA [56], CSNK2A1 interaction with NS1 plays an important part in parvovirus replication [57]; and RPL11 and RPS5 are components of the ribonucleoprotein complex [58,59]. Therefore, we hypothesize that these host factors may also be involved in the replication of RHDV.

**Figure 9.**
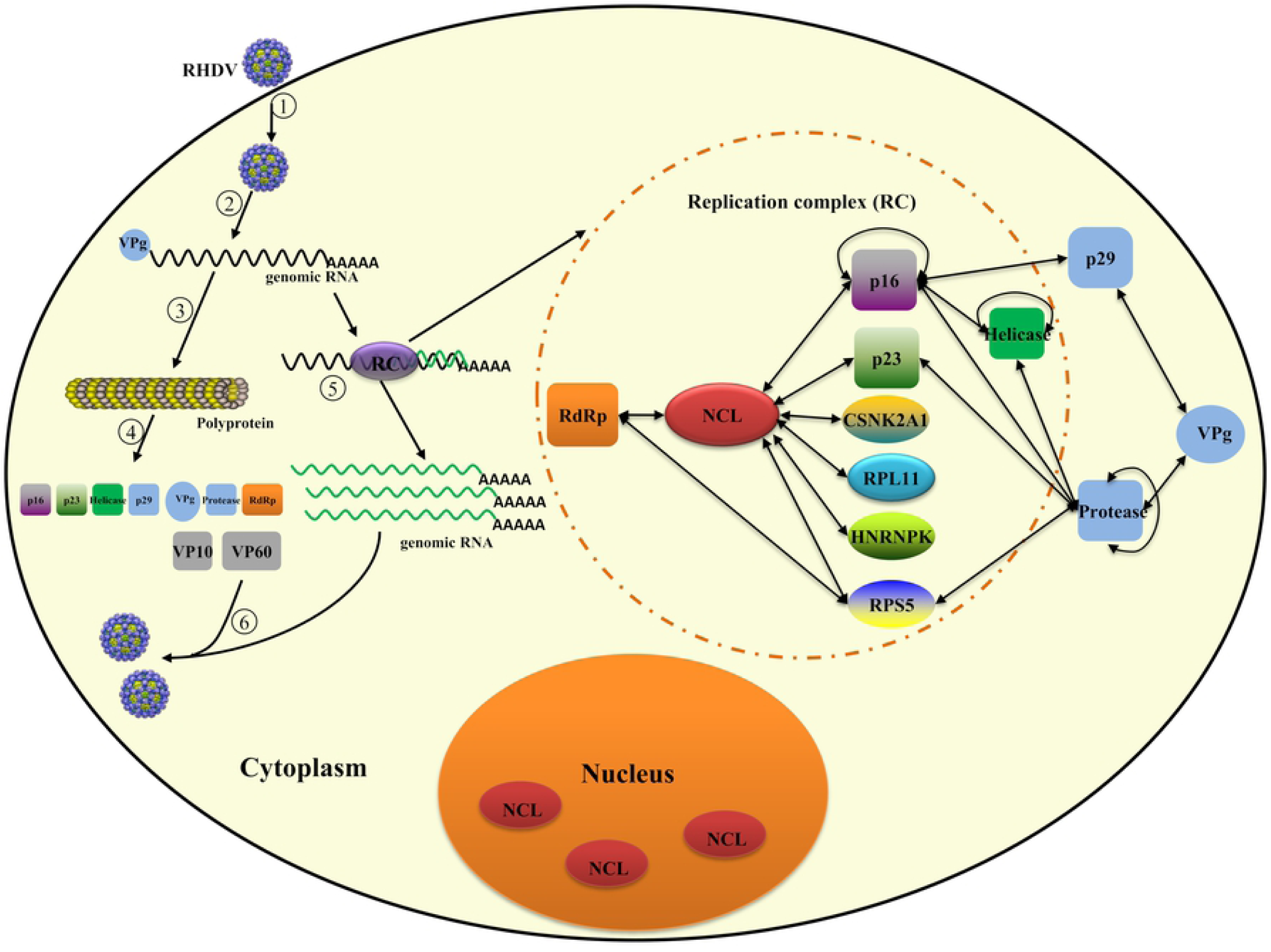
Schematic of the role of NCL in RHDV replication. (1) RHDV is internalized into the cell. (2) Uncoating of the viral genome. (3) Translation of the polyprotein precursor. (4) Polyproteins are cleaved into nonstructural and structural proteins. (5) RHDV genomic RNA is replicated in the RC. (6) Assembly of the structural proteins and genomic RNA. Components within the dashed line are the main components of the RHDV RC. The two-way arrow represents interaction between the two proteins.

We also identified that NCL interacts with nonstructural proteins (p16 and p23) of RHDV (Figs. 5, 6). By blocking the interaction of NCL with p16 and p23, we found that these interactions have important roles in RHDV replication (Fig. 7). Previous studies have reported that p16 can accumulate in subnuclear compartments, which may point to a specific interaction with nucleic acids and/or cellular proteins [42]. A similar nuclear/subnuclear accumulation of nonstructural proteins has been reported in picornavirus, for which nuclear accumulation of the 2A protein and a close association of this protein with the nucleolar ribosomal chaperone protein B23 have been reported. It was suggested that 2A upregulates the formation of modified ribosomes with a preference for viral internal ribosomal entry sites, thereby contributing to the inhibition of cap-dependent cellular mRNA translation [60]. Here, we revealed that NCL interacts with p16 of RHDV via the NTD domain (Fig. 5) and that this interaction plays a role in RHDV replication (Fig. 7). The N-terminal domain of NCL contains acidic regions, rich in glutamic acid and aspartic acid, which are the sites of phosphorylation and participate in the transcription of rRNA and interact with components of the pre-rRNA processing complex [24]. Therefore, we speculate that RHDV utilizes the interaction between p16 and NCL to hijack NCL-associated machinery of rRNA transcription and pre-rRNA processing to replicate viral gRNA. In addition, the role of p23 in RHDV replication is still unclear. Previous studies have found that, similar to other caliciviruses, RHDV p23 is enriched in the endoplasmic reticulum membrane and plays an important part in inducing intracellular membrane rearrangement [42]. Therefore, we believe that p23 interacts with NCL to recruit replication-associated host proteins to the endoplasmic reticulum membrane, and assembles these to form membrane-associated RCs.

NCL is an abundant and ubiquitously expressed protein in many growing eukaryotic cells [24]. NCL is mainly localized within the nucleolus, but it also exists in the nucleoplasm, cytoplasm, and cell surface [25,26]. NCL controls a wide range of fundamental cellular processes, such as ribosome biogenesis, proliferation, and cell cycle regulation, and it also has important roles in the infection process of multiple viruses [24]. For example, NCL acts as a receptor for human respiratory syncytial virus [61]. NCL also mediates cellular attachment and internalization of enterovirus 71 [62]. A recent study showed that NCL interacts with the capsid protein of dengue virus, suggesting a role in viral morphogenesis [63]. In previous studies, we also found that NCL mediates the internalization of RHDV through interacting with the RHDV capsid protein [64]. Of note, NCL also plays an important part in replication of several viruses. For example, NCL interacts with the FCV NS6 (protease) and NS7 (polymerase) proteins, and has a role in virus replication [34]. Similarly, the interaction between NCL and the UTRs of FCV [34] and poliovirus [65] stimulates translation of viral proteins. Moreover, NCL binds to a protein of herpes simplex virus type 1 to facilitate the export of US11 from the cell nucleus to the cytoplasm [66]. However, the function of NCL in binding to RHDV replicase and recruiting host factors to form RCs had not been revealed until now. Here, we demonstrate for the first time that NCL can specifically bind to RHDV replicase (RdRp) and can act as a link, recruiting host factors and viral proteins, to form RHDV RCs. Our findings enrich the current knowledge about the mechanism of NCL regulation of viral replication and provides new clues for further exploration of the interaction between RHDV and host proteins.

In conclusion, we identified the components of the RHDV RC, for the first time. We found that NCL acts as a link to recruit host factors, viral replicase (RdRp), and nonstructural proteins (p16 and p23), thereby forming a complex and ordered RHDV RC (Fig. 9). Elucidation of the molecular mechanism by which NCL regulates viral replicase assembly may lead to new insights into viral RC biogenesis and novel antiviral strategies.

## Materials and Methods

### Ethics Statement

All experiments were performed in a secondary biosecurity laboratory. All experiments involving rabbits were carried out in strict accordance with the recommendations of the Guide for the Care and Use of Laboratory Animals of the Ministry of Science and Technology of the People’s Republic of China, and all efforts were made to minimize suffering. All animal procedures were approved by the Institutional Animal Care and Use Committee of the Shanghai Veterinary Research Institute, Chinese Academy of Agricultural Sciences (permit number: SHVRIAU-18-035).

### Plasmids

The pRHDV-luc plasmid, in which the *VP60* and partial *VP10* genes are replaced with the *Fluc* gene, was generated in our previous study [14]. To generate pRHDV-luc-HA1, pRHDV-luc-HA2, pRHDV-luc-His1, and pRHDV-luc-His2, the nucleotide sequence encoding a hemagglutinin (HA) peptide (TAC CCA TAC GAT GTT CCA GAT TAC GCT) or a His_6_ peptide (CAT CAT CAT CAT CAT CAT) were inserted into the RdRp region using fusion polymerase chain reaction (PCR). The pRHDV-luc-His1/HA1, pRHDV-luc-His1/HA2, pRHDV-luc-His2/HA1, and pRHDV-luc-His2/HA2 plasmids, in which HA and His tags were simultaneously inserted in-frame into RdRp, were generated by fusion PCR (Fig. 1A). In addition, pRHDVΔp16-luc, pRHDVp16(^35^RHDF^38^)-luc, pRHDVΔp23-luc, pRHDVΔp23(90-145)-luc, pRHDVΔRdRp-luc, and pRHDVΔRdRp(448-478)-luc were constructed using fusion PCR.

The lentivirus-based expression plasmids were generated with a pLOV-CMV-GFP vector (Neuron Biotech, China) using In-Fusion HD Cloning kits (Clontech Laboratories, Inc., USA), according to the manufacturer’s instructions. The pLOV-CMV-GFP vector was linearized with N*he* I and N*ot* I. The packaging plasmid psPAX2 containing the specific lentivirus genes Gag, Pol, and so on, and the pMD2.G plasmid, which expresses the G protein of the vesicular stomatitis virus (VSV-G), were purchased from Neuron Biotech.

The plasmids, used in M2H assays, were generated with the pACT and pBIND vectors (Promega Corporation, USA) using In-Fusion HD Cloning kits. The pG5luc vector (Promega) contains five GAL4 binding sites upstream of the TATA box, which controls *Fluc* expression. The pGL4.75 vector (Promega) encodes the luciferase reporter gene *Renilla reniformis* (*Rluc*) from a CMV promoter.

The p3×FLAG-CMV-14 vector (Sigma-Aldrich Corporation, USA) and pCMV-Myc (Clontech Laboratories, Inc.) were used to create mammalian expression constructs. The pGEX-4T-1 vector (GE Healthcare Life Sciences, USA) was expressed in competent *E. coli* BL21-CodonPlus (DE3) cells. *NCL* (GenBank accession number XM_017343189.1), *CSNK2A1* (GenBank accession number NM_001160284.1), *HnRNPK* (GenBank accession number NM_001082125.1), *RPL11* (GenBank accession number XM_008265690.2), and *RPS5* (GenBank accession number XM_002721896.3) sequences were amplified by reverse transcription PCR (RT-PCR) from an RK-13 cell cDNA library. Total RNA was isolated from RK-13 cells using TRIzol reagent (Invitrogen Corporation, USA), according to the manufacturer’s instructions. DNA was removed from the isolated RNA using DNase I (Takara Bio, Inc., Japan) and cDNA was produced with Moloney murine leukemia virus reverse transcriptase (M-MLV RT) (Promega) and random hexamers. RHDV genes (*p16, p23, helicase, p29, VPg, protease*, and *RdRp*) were amplified using RT-PCR from RHDV cDNA. The genomic sequence of RHDV CHA/JX/97 was retrieved from the GenBank database (accession number DQ205345). Viral cDNA was generated as described in our previous report. All plasmids were created using In-Fusion HD Cloning kits, according to the manufacturer’s instructions.

All RT-PCR and PCR amplifications for cloning were performed with TransStart® FastPfu Fly DNA Polymerase (TransGen Biotech Co., Ltd., China), according to the manufacturer’s instructions. RT-PCR and PCR products were separated by agarose gel electrophoresis and purified with a SanPrep Column DNA Gel Extraction Kit (Sangon Biotech Co., Ltd., China). Restriction digests were performed using commercial kits (New England Biolabs, USA), according to the manufacturer’s instructions. All plasmid sequences were amplified by PCR and analyzed by Sanger sequencing, to verify the sequence fidelity and reading frames (Sangon Biotech). The details of all constructs used in the study, including residue numbers, expression vectors, and tags, are summarized in Table S1. In addition, the primers used in this research are listed in Table S2.

### Cell lines and viruses

Rabbit kidney cells (RK-13, ATCC, CCL37) and 293T cells (ATCC, CRL-3216) were routinely maintained in minimal essential medium (MEM) (Life Technologies, USA) or Dulbecco’s modified Eagle’s medium (DMEM) (Life Technologies), respectively, supplemented with 10% fetal bovine serum (Biological Industries, Israel). RK-NCL cells, RK-NCL-NTD cells, RK-NCL-RBD cells, RK-NCL-CTD cells, and RK-GFP cells were generated by transducing RK-13 cells with VSV-G pseudotyped lentiviral particles, which contain the *NCL* gene, NCL-NTD domain, NCL-RBD domain, NCL-CTD domain, or *GFP* gene. To generate those recombinant lentiviral particles, we transfected psPAX2 (10 μg), pMD2.G (12 μg), and pLOV-NCL-GFP (22 μg), pLOV-NTD-GFP (22 μg), pLOV-RBD-GFP (22 μg), pLOV-CTD-GFP (22 μg) or pLOV-CMV-GFP (22 μg), respectively, into 293T cells, which were seeded (1×10^6^ cells) onto 100-mm tissue culture dishes, using a calcium phosphate transfection reagent (Invitrogen). The lentivirus packaging and transduction procedures are based on our previous studies [4]. For the RHDV replicon replication assay, these five stable cell lines, with exactly the same number of passages, were used to avoid the effect of cell passage variation on RHDV replication.

RHDV strain JX/CHA/97 was isolated in 1997 during an outbreak of RHDV in China and stored in our laboratory. The genomic sequence of RHDV CHA/JX/97 is available in the GenBank database (accession number DQ205345). mRHDV was mutated from RHDV strain JX/CHA/97 and stored in our laboratory [15].

### Antibodies and chemicals

The antibodies used in this study included: mouse anti-His, anti-HA, anti-luc, and anti-myc antibodies purchased from Abcam; mouse anti-Flag obtained from Sigma-Aldrich; mouse anti-β-actin and anti-GST antibodies purchased from Kangwei Century Biotechnology; mouse anti-NCL and rabbit anti-NCL obtained from Thermo Fisher Scientific; mouse anti-RHDV RdRp was prepared by Genscript and stored in our laboratory; polyclonal rabbit anti-RHDV p16 and anti-RHDV p23 were prepared and stored in our laboratory; goat anti-mouse IgG conjugated with HRP and goat anti-rabbit IgG conjugated with HRP purchased from Jackson ImmunoResearch Europe Ltd.; goat anti-mouse IgG conjugated with Alexa Fluor 488, goat anti-rabbit IgG conjugated with Alexa Fluor 488, goat anti-mouse IgG conjugated with Alexa Fluor 633, and goat anti-rabbit IgG conjugated with Alexa Fluor 633, obtained from Thermo Fisher Scientific. DAPI staining solution purchased from Beyotime Biotechnology. N*he* I and N*ot* I obtained from New England Biolabs. CCK-8 kit purchased from Dojindo Laboratories. RdRp peptide (ERGVQLEELQVAAAAHGQEFFNFVRKELER), p16 peptide (PLSFFLDLRDKTPPCCIRAT) and non-specific peptide (HKFGPVCLCNRAYIHDCGRW) were synthesized by GL Biochem (Shanghai) Ltd, and the purity was 90%. We dissolved these peptides in PBS at a concentration of 10 ng/μL. Then, these peptides were directly added into the cell culture medium at different working concentrations, respectively, and these peptides entered in the cytoplasm by macropinocytosis.

### Affinity purification of protein complex

RK-13 cells were seeded onto ten 100-mm tissue culture dishes at a density of 1× 10^6^ cells/dish. The cells were grown overnight and then transfected with pRHDV-luc-His1/HA1 (12 μg /dish) using Lipofectamine 3000 (Invitrogen), according to the manufacturer’s instructions. After 48 hpt, the cells were washed three times with cold Modified Dulbecco’s phosphate-buffered saline (MDPBS; Thermo Scientific™ Pierce™, USA). The cells were then lysed with 500 μL/dish of ice-cold lysis buffer (Tris, 0.15 M NaCl, 0.001 M EDTA, 1% NP-40, 5% glycerol; pH 7.4, proteinase inhibitor cocktail (Thermo Scientific). After incubating cells on ice for 5 min with periodic mixing, the lysate was transferred to a microcentrifuge tube and centrifuged at ∼ 13,000 × g for 10 min to pellet the cell debris. The soluble fraction was incubated by head-to-tail rotation with 300 μL of anti-HA antibody-coated beads (Thermo Scientific) for 4 h at 4°C. The beads were collected by centrifugation and then washed four times with 10 mL washing buffer (Tris, 0.15 M NaCl, 0.001 M EDTA, 1% NP-40, 5% glycerol; pH 7.4, 20 mM imidazole (Thermo Scientific). After being washed, the bound proteins were eluted with 400 μL of wash buffer supplemented with 250 μg/mL HA peptide (Sigma-Aldrich) by incubation at room temperature for 10 min. After centrifugation at 3,000 × g for 5 min, followed by a second elution with 200 μL of wash buffer supplemented with 250 μg/mL HA peptide. The eluted solutions were combined and diluted into 1.5 mL of lysis buffer containing 20 mM imidazole, and 60 μL of Ni Sepharose (Thermo Scientific) was added. After incubation at 4°C for 1 h and clarification, the beads were washed four times with 1.5 mL wash buffer. The captured proteins were eluted in 80 μL wash buffer containing 240 mM imidazole and mixed with 20 μL of 5X SDS loading buffer (250 mM Tris Cl (pH 6.8), 30% glycerol, 10% SDS, 0.02% bromophenol blue, 25% 2-mercaptoethanol). After being boiled for 10 min, the proteins samples were separated by SDS-PAGE, and protein bands were visualized with silver staining.

### Quantitative reverse transcription (qRT)-PCR

Total RNAs were purified using TRIzol reagent (Invitrogen), according to the manufacturer’s instructions. DNA was removed from the isolated RNA using DNase I (Takara), and then cDNA was produced using M-MLV RT (Promega) and random hexamers. The cDNA samples were subjected to real-time PCR with SYBR Premix *Ex Taq* Tli RNase H Plus (Takara) using an ABI 7500 Fast Real-Time PCR system (Applied Biosystems, USA). The primers are listed in Table S2. The relative RNA levels were determined according to the 2^− ΔΔCT^ method. The amount of mRNA in each sample was normalized to that of GAPDH.

### Mammalian two-hybrid (M2H) assays

The interactions between host protein and RHDV nonstructural proteins were evaluated using a CheckMate Mammalian Two-Hybrid System (Promega). The proteins expressed from the pACT vector recombinant plasmid acted as prey proteins, and proteins expressed from the pBIND vector recombinant plasmid acted as bait proteins. Subsequent M2H analysis was performed, according to the manufacturer’s instructions. In brief, bait and prey plasmids were co-transfected with pG5luc plasmids into subconfluent 293T cells at a molar ratio of 1:1:1 for pACT: pBIND: pG5luc vector. At 48 h after transfection, the 293T cells were lysed, and Renilla luciferase (Rluc) and firefly luciferase (Fluc) activities were evaluated using the Dual-Luciferase Reporter (DLR™) Assay System (Promega).

### Luciferase activity measurements

Cells were washed with PBS and lysed in 200 μL of 1X Passive Lysis Buffer (Promega). After gentle shaking for 15 min at room temperature, the cell lysate was transferred to a tube and centrifuged for 2 min at 12,000 × g at 4°C. The supernatant (20 μL) was added to 100 μL of luciferase assay substrate to evaluate the activity of Fluc and Rluc using the Promega DLR™ assay system, based on relative light units (RLUs). Luciferase activity was analyzed using a FB12 luminometer (Berthold, Germany). To normalize the luciferase values determined for cells transfected with the Fluc replicon, Rluc activity was used as an internal control.

### Bacterial expression of recombinant proteins and purification

All proteins were expressed in competent *E. coli* BL21-CodonPlus (DE3) cells (TransGen Biotech Co., Ltd.) that were seeded in 1 mL of an overnight starter culture and then grown in 100 mL of Luria-Bertani (LB) broth with shaking at 220 rpm at 37°C to mid-log phase (∼0.6–0.8 OD_600_). Cells were then typically induced with 0.2–0.5 mM isopropyl β-D-1-thiogalactopyranoside and incubated for approximately 16 h at 16°C with shaking at 220 rpm. The details of protein expression are available on request. Cells were pelleted by centrifugation at ∼5,000 × g and stored at -80°C. Bacterial pellets were resuspended in lysis buffer (20 mM Tris/HCl; pH 7.4, 60 mM NaCl, 1 mM ethylenediaminetetraacetic acid (EDTA), 1 mg/mL lysozyme, 1 mM dithiothreitol, and 0.1% Triton X-100) supplemented with complete protease inhibitor cocktail (Thermo Scientific) for 1 h on ice. Nuclease was then added and the lysate was incubated for 1 h at ambient temperature under rotation. The lysates were centrifuged at 4°C for 10 min at 12,000 × g. Glutathione Sepharose 4B beads (Pierce Biotechnology, USA) were added to the clarified supernatants and the mixtures were incubated overnight at 4°C under rotation. The beads were washed with lysis buffer, followed by three washes with PBS, and then stored at 4°C in an equal volume of PBS.

### Glutathione-S-transferase (GST) pull-down assay

For the *in vitro* binding assay, Flag- or myc-tagged NCL, and RHDV p16, p23, and RdRp proteins were expressed in RK-13 cells. According to the manufacturer’s instructions, the GST pull-down assay was performed by incubating 50 μL of a 50% slurry of glutathione Sepharose beads containing 25 μM GST fusion protein in lysis buffer with a 3-fold molar excess of prey protein (Pierce Biotechnology). RNase (Takara) was added to the cell lysis and wash buffers. The bound proteins were separated by SDS-PAGE and then subjected to western blot analysis.

### Co-immunoprecipitation (Co-IP) analysis

RK-13 cells were co-transfected with the bait and prey plasmids. At 48 h after transfection, total protein was isolated from RK-13 cells using IP lysis buffer. We conducted Co-IP analysis using a commercial Co-IP kit (Pierce Biotechnology), according to the manufacturer’s instructions. AminoLink Plus Coupling Resin (Thermo Scientific) was incubated with anti-myc monoclonal antibody (mAb) (Abcam, UK) or anti-Flag mAb (Abcam) and then subjected to SDS-PAGE. IB analysis of the proteins was subsequently conducted using mAbs against myc and Flag (Abcam). RNase was added to the cell lysis and wash buffers.

### Immunoblotting (IB) analysis

Protein samples were separated on 12% gels and then transferred to nitrocellulose membranes (Hybond-C; Amersham Life Sciences, UK) using a semi-dry transfer apparatus (Bio-Rad Laboratories, USA). The membranes were blocked with 5% (w/v) nonfat milk in TBST buffer (150 mM NaCl, 20 mM Tris, and 0.1% Tween-20; pH 7.6) for 3 h at 4°C and then stained overnight at 4°C with a primary antibody (Ab). After washing three times for 10 min each, the membranes were incubated with a secondary Ab against immunoglobulin G (IgG) conjugated to horseradish peroxidase (Sigma-Aldrich) in PBST buffer (137 mM NaCl, 2.7 mM KCl, 10 mM Na_2_HPO_4_, 2 mM KH_2_PO_4_ and 0.1% Tween-20; pH 7.4) for 1 h at room temperature (RT). Finally, after washing three times for 10 min each, the proteins were detected using an automatic chemiluminescence imaging analysis system (Tanon Science & Technology Co., Ltd., China).

### Immunofluorescence assay (IFA)

Cells were fixed in 3.7% paraformaldehyde in PBS (pH 7.5) at RT for 30 min and subsequently permeabilized by incubation in methanol at -20°C for 30 min. The fixed cells were blocked with 5% (w/v) nonfat milk in PBST buffer for 3 h at 4°C and then stained with a primary Ab for 2 h at 37°C. After washing three times for 10 min each, the cells were incubated with a secondary Ab against IgG conjugated to fluorescein isothiocyanate (FITC) (Sigma-Aldrich) in PBST buffer for 1 h at room temperature. Finally, after washing three times for 10 min each, the samples were observed under a fluorescence microscope equipped with a video documentation system (ZEISS, Germany).

### Mass spectrometry

Jingjie PTM BioLab Co., Ltd. (Hangzhou, China) performed all mass spectrometry analyses.

### In vivo experiments

Twenty 8-week-old male New Zealand rabbits seronegative for RHDV were randomly distributed into four groups (n = 5/group) and housed in individual ventilated cages. All experimental protocols were reviewed by the state ethics commission and were approved by the competent authority. Details of the protection assay are shown in Fig. 7F. All rabbits were challenged intramuscularly with 100 × the median lethal dose (LD50) of RHDV. Two hours after RHDV infection, rabbits were injected with the RdRp peptide (1 mg), p16 peptide (1 mg), non-specific peptide (1 mg), or PBS, via the ear vein. The rabbits were clinically examined daily for 7 days post-challenge.

### Statistical analyses

Statistical analysis was performed using GraphPad Prism 6 software. Specific tests are described in the figure legends.

## Acknowledgments

We thank LetPub for its linguistic assistance during the preparation of this manuscript.

## Supporting information

**S1 Fig. RHDV RdRp protein structure analysis.** For clarity, the structure of the RHDV RdRp was obtained from the Protein Data Bank under the identification number 1khv (https://www.rcsb.org/). The structure of the RdRp mutation, as predicted by the SWISS-MODEL online tool (https://www.swissmodel.expasy.org/) and based on homology molecules found in the Protein Data Bank. The orange portion indicated by the red arrow is the inserted label.

**S2 Fig. Effect of NCL siRNA on RK-13 cells. (A)** After RK-13 cells were transfected with different amounts of NCL siRNA for 24 h, the number and activity of viable cells were detected using CCK-8. **(B)** In different doses of NCL siRNA-treated RK-13 cells, we transfected the RHDV replicon reference plasmid pRluc. After 24 hpt, cell lysates were collected and the activity of Rluc was detected using a dual luciferase reporter system. Student t-tests and analysis of variance were used for statistical analyses. *p < 0.05 and **p < 0.01. The number of cells used in all replicate experiments was similar.

**S3 Fig. Alignments of amino acid sequences of G1.1a–G1.2 genogroups of RHDV. (A)** Alignments of amino acid sequences 448–478 of RdRp protein of G1.1a–G1.2 genogroups of RHDV. The representative strains used in the alignments are G1.1a: RHDV strain JX/CHA/97 (GenBank accession no. DQ205345.1) and RHDV isolate K5_08Q712_BatchRelease1/2008 (GenBank accession no. MF598301); G1.1b: RHDV strain CB194_Pt (GenBank accession no. JX886001) and RHDV-SD (GenBank accession no. Z29514.1); G1.1c: RHDV isolate BlueGums-2 (GenBank accession no. KT280058) and RHDV isolate AUS/NSW/OUR-1/2014/06 (GenBank accession no. KY628318); G1.1d: RHDV-FRG (GenBank accession no. M67473) and RHDV-FRG/2000 (GenBank accession no. NC001543); G1.2: RHDV isolate CBAlgarve14-1 (GenBank accession no. KM115714) and RHDV isolate BLMT-1 (GenBank accession no. KT280060). Sequence alignment was performed with the ClustalW algorithm (http://www.clustal.org/). The conserved NCL-binding motif is boxed. Selected amino acids from this motif are indicated with RHDV RdRp numbering. **(B)** Alignments of representative amino acid sequences of the p16 protein of G1.1a–G1.2 genogroups of RHDV. Sequence alignment was performed with the ClustalW algorithm. The conserved NCL-binding motif is boxed. Selected amino acids from this motif are indicated with RHDV p16 numbering. **(C)** Alignments of representative amino acid sequences of p23 protein of G1.1a–G1.2 genogroups of RHDV. Sequence alignment was performed with the ClustalW algorithm. The conserved NCL-binding motif is boxed. Selected amino acids from this motif are indicated with RHDV p23 numbering.

**S4 Fig. Expression level of the domain of NCL in overexpression cells. (A)** The positive rate of cells was observed under a fluorescence microscope. GFP-fused NCL domain (NTD, RBD, or CTD) was expressed in RK13-NCL-NTD, RK13-NCL-RBD, and RK13-NCL-CTD cells, respectively. **(B)** The expression level of the domains of NCL in the overexpression cells at 10 passages were determined by western blot analysis with anti-Flag mAb. RK13-GFP cells were used as positive control. β-actin was used as an internal control.

**S1 Table. Plasmid construct details.**

(XLSX)

**S2 Table. Oligonucleotide primer sequences.**

(XLSX)

**S3 Table. Details of protein expression.**

(XLSX)

